# Social bond dynamics and the evolution of helping

**DOI:** 10.1101/2023.10.11.561838

**Authors:** Olof Leimar, Redouna Bshary

**Affiliations:** Department of Zoology, Stockholm University, 106 91 Stockholm, Sweden; Institute of Biology, University of Neuchâtel, Neuchâtel, Switzerland

**Keywords:** Reciprocity, learning, interdependence, game theory

## Abstract

Empiricists often struggle to apply game theory models to real-life cases of animal cooperation. One reason is that many examples of cooperation occur in stable groups, where costs and benefits of helping depend on several factors. Among these are variable investments, fitness interdependencies, learning, memory, reciprocity, and partner choice, including the formation of social bonds with specific group members. Here, we present a game theory model exploring the conditions under which social bonds between group members can promote cooperation, with reciprocal acts of helping. In the model, bonds build up from exchanges of help in a similar way as the strength of association increases in learning, as in the Rescorla-Wagner rule. The bonds in turn affect partner choice and influence helping amounts. The model has a mechanism of reciprocity for bonded pairs, which can evolve towards either loose or strict reciprocation. Several aspects of the model are inspired by observations of food sharing in vampire bats. We find that small social neighbourhoods are required for the evolutionary stability of helping, either as small group sizes, or if members of larger groups can form temporary (daily) smaller groupings. The evolutionary outcome is a fairly low cost helping, while the benefit of receiving help can be substantial. Individuals in need request help based on bond strength, but there is also an evolved preference for initiating bonds with new group members. In contrast, if different groups come into temporary contact with each other, the evolved tendency is to avoid forming bonds between groups.

**Popular science summary:** The search for evolutionary explanations of cooperation between members of social groups has long been a high profile endeavour. A case of particular interest is when individuals develop a network of friends and exchange help through these social bonds. The question of helping between friends was given emphasis already from the start of the evolutionary study of cooperation, more than 50 years ago, but it has remained without any decisive progress since that time. Here we present a game-theory analysis of helping through the build-up of social bonds. We find that there is reciprocity in socially bonded pairs, which is neither immediate nor very strict, and that relatively small social neighbourhoods are required for the evolutionary stability of helping.

---

Over a period of several decades, game-theory modelling has made important contributions to the study of the evolution of cooperation [1]. The modelling has shown that strategies of investment into a partner can be evolutionarily stable and result in lifetime direct fitness benefits, verifying in principle that cooperation could evolve. Still, a remaining and even more challenging task is to apply game-theory models to real-life cases of cooperation. A number of examples involve species that live in fairly stable groups, where individuals seldom change group after becoming adult. The individuals typically do not interact randomly with other group members, but tend instead to have one or a few preferred partners. These enduring affiliative relationships are described as social bonds [2, 3]. Bonded pairs spend more time in proximity to each other, have more favourable interactions, such as grooming, coalition formation and food sharing, and are generally more tolerant towards each other. The evolution of helping through social bond formation is thus a biologically relevant example of cooperation, but models that explain when this form of helping can evolve are currently lacking. Here, we provide such a model.

There are several challenges facing the modelling, including accounting for individual variation and specifying the cognitive and behavioural mechanisms of bond formation. The bonds should influence decisions about who to request help from and how much help to provide to different partners, while accounting for the possibility of providing no or very little help. We assume that social bonds build up during exchanges of help in a similar way as the strength of association in Pavlovian learning. In our model, individuals have genetically determined bond build-up or learning rates, which can evolve. If bonds build up, individuals in need can use bond strengths to decide who to ask help from, using a choice mechanism with bond strengths as values. Bond strengths can also be used when individuals in a group form temporary (daily) subgroups, which allows for closer association between bonded individuals, provided that this tendency to associate evolves. The tendency (positive or negative) to initiate bonds with new group members, the maximum amount of help provided, and the degree of enforcement of reciprocity within a bond are additional model elements. We examine the evolution of genetically determined traits that influence these different aspects of the helping interactions, using individual-based simulation of large populations of haploid, sexually reproducing individuals.

In our model analyses, we first study how helping with social bonds might operate, in terms of the build-up of bond strength, the helping amounts, the extent of reciprocity, and the degree of association between partners. We then investigate if there are requirements on the sizes of social neighbourhoods (groups and sub-groupings) for this type helping to be evolutionarily stable. We also investigate how readily new social bonds are initiated, and whether this depends on the type of contacts between individuals, either as new permanent group members, or as temporary contacts between members of different groups. To avoid any effects of relatedness on helping, new recruits into a group are derived from parents chosen from the entire population.

Survival is the fitness component acting in the model, with helping in the form of food sharing that leads to survival costs and benefits for donor and recipient individuals. In this way, individuals in bonded pairs, as well as the members of small groups, can become dependent on each other for survival. The model is inspired by observations of food sharing in vampire bats [4], for which there are data on the extent of reciprocity [5, 6].

After presenting the model and results, we discuss the similarities and differences between our analysis here and previous game-theory-inspired ideas about helping in groups [7, 8, 9, 10, 11, 12], including the concept of raising-the-stakes cooperation [13, 14] and the formation of social relationships [15]. We emphasize the potential value of implementing sufficient details of the cognitive and behavioural mechanisms operating in real-life examples when attempting to explain the evolution of cooperation.

## Model and results

Here we give an overview of the model, with a detailed description in the Methods section and the Supplementary Information. Individuals spend their lives in groups of size *N*, and time is divided into steps, where a time step might be one day. A group can be structured into temporary subgroups, which we identify with places, for instance resting places (for vampire bats, a resting place might be a daytime roost [4]). There are *K* places for a group, with on average *G* = *N/K* individuals per place, and each day each group member can select which place to go to. If two individuals go to the same place, they can help each other.

We use a state variable *z*_*i*_ to indicate the current (nightly) foraging success of individual *i*. The state is *z*_*i*_ = 1 with probability *p*_s*i*_ (the individual succeeded in foraging) and *z*_*i*_ = 0 with probability 1 *− p*_s*i*_. An individual’s probability of foraging success depends on its phenotypic quality *q*_*i*_. Only individuals with *z*_*i*_ = 0 ask for help on a given day. They only ask help from other individuals *j* they perceive have a state of *z*_*j*_ = 1, and they have a probability *p*_d_ of correctly detecting that state.

### Bond strength

We use *x* and *y* to denote the strength of bonds. For an individual *i* who is in need of help on a given day (i.e., *z*_*i*_ = 0), there is an updating of bond strengths. The starting value of the bond strength is *x*_s_, which should be large enough that at least some help is donated early in a partnership. After a request from *i* to an individual *j, i* updates its value *x*_*ij*_ using a genetically determined learning rate *α*_*i*_ and an asymptotic (maximum) bond strength *x*_a_. If *u*_*ji*_ is the amount of help from *j* to *i*, the update is

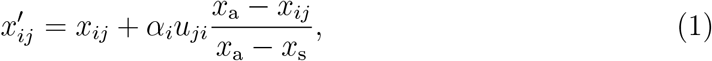

where 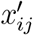 is the updated value of *x*_*ij*_. The donating individual *j* also updates its bond strength *x*_*ji*_, as follows:

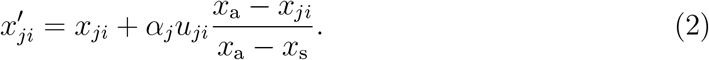

By having both *i* and *j* update their bond strength values, there is approximate symmetry in the bond dynamics. This bond strength dynamics is inspired by the Rescorla-Wagner model of classical conditioning [16] and is illustrated in Fig. 1a.

**Figure 1:**
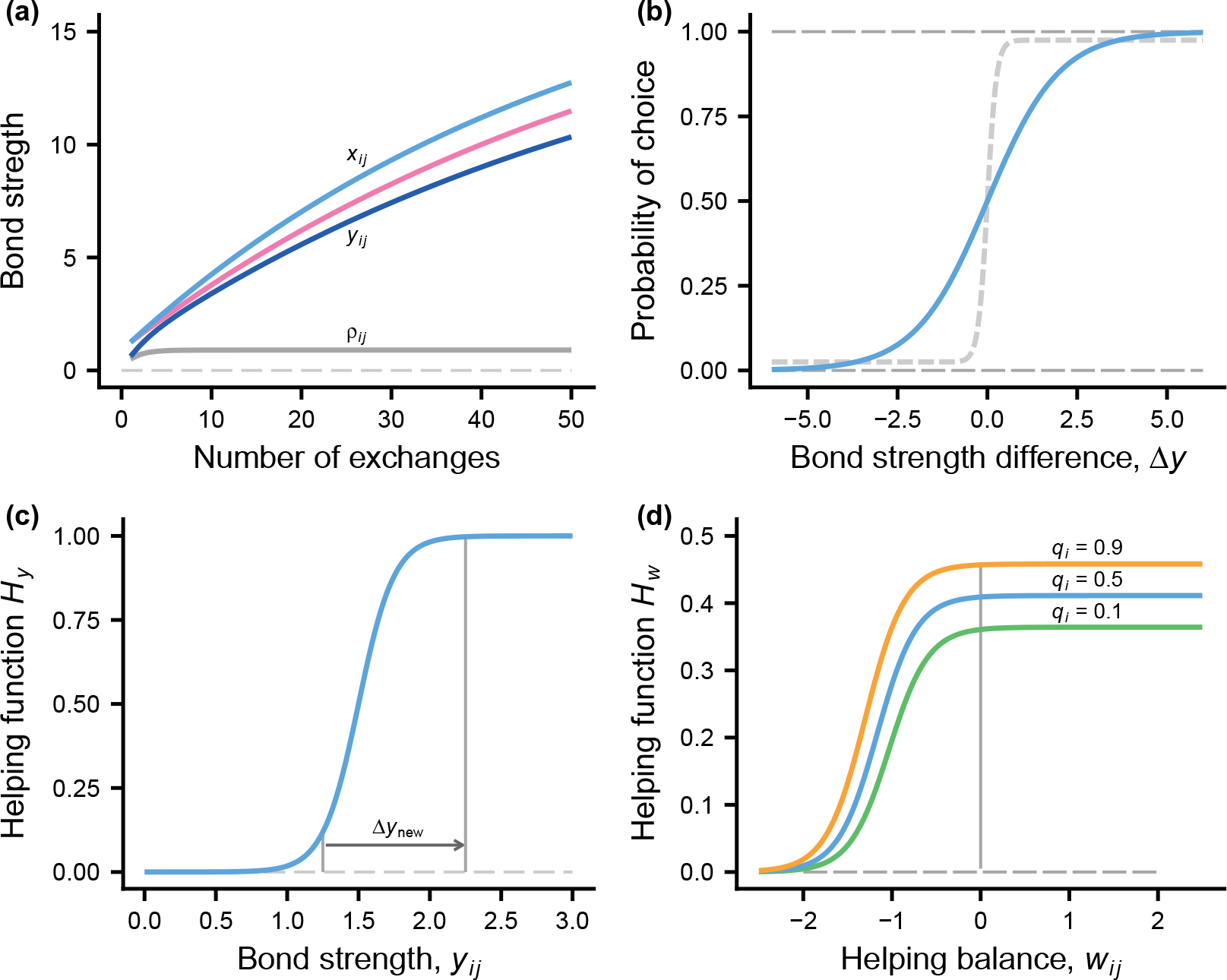
Model elements. **(a)** Stylised bond strength dynamics. In the model, the bond strength *x*_*ij*_ between individuals *i* and *j*, as perceived by *i*, increases with each helping exchange. Individuals have genetically determined learning rates *α*_*i*_ and *α*_*j*_ which, together with the helping amounts *u*, determine the bond strength increments. The light blue and red curves show *x*_*ij*_ and *x*_*ji*_, assuming *α*_*i*_ = 1.2 and *α*_*j*_ = 1.0 and all helping amounts *u* = 0.3. The dark blue curve shows the effective bond strength *y*_*ij*_, which is *x*_*ij*_ multiplied by an estimate *ρ*_*ij*_ of association between *i* and *j* (0 *≤ ρ*_*ij*_ *≤* 1), which indicates how often *i* and *j* have been in the same place in the recent past. **(b)** Soft-max probabilities of choice by an individual between two alternatives, as a function of the difference ∆*y* in effective bond strength. The blue/dashed light grey curves give the choice between two partners/places. **(c)** The dependence of the helping function *H*_*y*_ on the effective bond strength *y*_*ij*_ = *ρ*_*ij*_*x*_*ij*_. The arrow illustrates a possible extra effect of interacting with a new partner. **(d)** The dependence of the helping function *H*_*w*_ on the helping balance *w*_*ij*_ between individuals *i* and *j* (the curves are for *z*_*i*_ = 1 and different values of *q*_*i*_). The amount *u* of help donated is the product *u* = *H*_*w*_*H*_*y*_ of the two functions.

To account for the recent same-place association between individuals, we introduce an effective (association adjusted) bond strength *y*_*ij*_, defined as the product of an estimated association *ρ*_*ij*_ and the bond strength *x*_*ij*_, the latter representing the accumulated exchange of help between *i* and *j*. So, for *i* the effective strength of the bond to *j* is *y*_*ij*_ = *ρ*_*ij*_*x*_*ij*_. This effective bond strength can decrease over time, which happens if previously bonded individuals are mostly in different places.

For an individual in need, if there is more than one other individual to ask help from, the effective bond strengths are used as values for a soft-max choice, which is illustrated in Fig. 1b for a choice between two alternatives. Evolutionary changes in the learning rates *α*_*i*_ will influence such differences in bond strength, so this partner choice mechanism can be modified by evolution. Furthermore, the daily choices of place to go to are influenced by an individual’s recent experience of the average effective bond strengths to other individuals encountered in different places. To study the evolution of the choice of place, we assume that individuals have a genetically determined trait *β*_*i*_ that influences their sensitivity to place differences in estimated average bond strength (*β*_*i*_ can be positive, zero, or negative). An illustration of the resulting probability of choice between two places appears in Fig. 1b.

An individual in need can ask for help from more than one of the available group members (one reason for introducing this is that it seems to occur in vampire bats). In the model, individuals in need have the opportunity to ask for help twice, from different individuals among the available ones in the current place.

### Determination of helping amounts

The amount *u*_*ij*_ donated by *i* when *j* asks for help depends of several factors. First, there is a sigmoid dependence on the effective bond strength *y*_*ij*_, which is shown as the helping function *H*_*y*_ in Fig. 1c. This form of dependence allows for successive increases in the helping amounts, as in raise-the-stakes cooperation. To investigate evolutionary modifications in how new individuals are treated, individuals have a genetically determined trait ∆*y*_new *i*_, which acts as a temporary increment to the effective bond strength on the first occasion of donating to an individual, illustrated by the arrow in Fig. 1c. This increment can evolve to be positive, zero, or negative. The increment is also applied to new individuals when choosing who to ask help from (Fig. 1b).

The model has another helping function *H*_*w*_, illustrated in Fig. 1d. The amount of help donated by *i* to *j* is the product of the two functions, *u*_*ij*_ = *H*_*y*_*H*_*w*_. The function *H*_*w*_ determines the maximum amount of help and the extent of reciprocity, through two genetically determined traits *h*_a*i*_ and *h*_s*i*_, with *h*_a*i*_ being the maximum amount of help possible from a highest quality individual (*q*_*i*_ = 1) and *h*_s*i*_ determining the sensitivity to the total helping balance *w*_*ij*_. The total helping balance is the total previously received minus donated for *i* interacting with *j*. Provided that *h*_s*i*_ evolves to a positive value, it follows that if *w*_*ij*_ becomes too negative, the amount *i* donates to *j* becomes small and approaches zero (Fig. 1d). Further details about the helping functions are given in Methods.

### Mortality

The day-to-day mortality depends on an individual’s foraging success (*z*_*i*_), but also on its resource balance *ζ*_*i*_ for a given day (Fig. 2a). The current day resource balance increases when an individual receives help and decreases when it donates. At the start of a day, *ζ*_*i*_ is assumed to depend on the phenotypic quality *q*_*i*_, such that a higher quality individual suffers a smaller mortality increase when donating a certain amount. A consequence of the dependence illustrated in Fig. 2a is that the survival cost of donating starts out small but accelerates for larger amounts, whereas for receiving there is first a sharp decrease in mortality, which flattens out for larger amounts.

**Figure 2:**
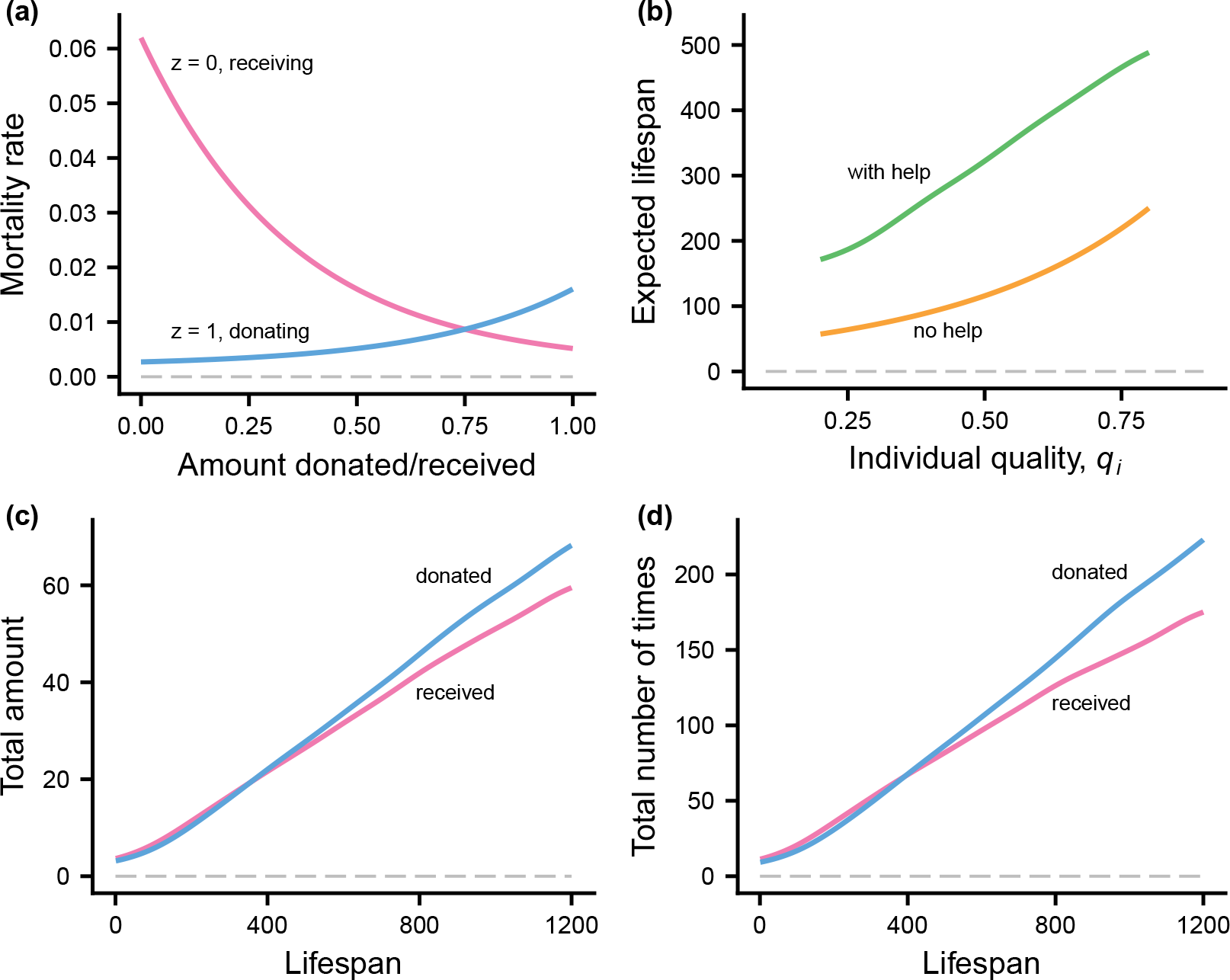
Mortality and lifespan in the model. **(a)** Day-to-day mortality for successful foragers (*z* = 1) that donate help (blue) and unsuccessful foragers (*z* = 0) that receive help (red), as a function of the amount transferred (i.e., the change in resource balance *ζ*). **(b)** Effects of individual phenotypic quality *q*_*i*_ on expected life span, with and without helping. **(c)** Average total amounts of help donated (blue) and received (red) by an individual, as a function of the individual’s lifespan. **(d)** Average total number of times help was donated (blue) and received (red) by an individual. The curves are kernel smoothing fits and are based on ca 12 000 individuals with complete life histories. There is considerable variation around the fitted curves (see Fig. S1). The data for panels **(b)** to **(d)** come from a simulation with 175 groups of size *N* = 24, where each group has access to *K* = 6 places (*G* = 4). The simulation was run at evolutionary equilibrium for 4000 days, and the data come from individuals born in the earlier part of this interval, thus including those with very long lifespans.

Individuals that die are replaced through reproduction by surviving individuals. Replacement happens at intervals of *T* periods (we used *T* = 20). To avoid effects of local relatedness, parents to new individuals are randomly drawn from the global population.

### Examples of helping with social bonds

To illustrate helping in the model, we present evolutionary equilibrium results for a case with groups of size *N* = 24 and *K* = 6 places (*G* = 4). Figure 2b shows that helping substantially increases the expected lifespan compared to a hypothetical case where there is no helping, with a proportionally greater increase for lower quality individuals.

The average number of times and total amounts for donating and receiving help increase approximately in proportion to life span (Fig. 2c, d). Higher quality individuals tend to live longer and on average have relationships where the partner needs help more than they themselves do, which explains the divergence between donated and received help for long lifespans. In the simulation, an average quality individual needs help on one out of every ten days.

An example of the exchange of help between two individuals appears in Fig. 3, coming from the same simulation as in Fig. 2. The build-up of bond strength in Fig. 3a is less smooth than for the stylised case in Fig. 1a. The reason is that the helping balance *w* varies (Fig. 3b), in this example in such a way that one of the individuals mostly has a negative balance (i.e., donating more than receiving; blue curve), causing the individual to reduce the amount it donates when the other asks for help (Fig. 3c). The individual with mostly negative helping balance still donates a larger total amount than it receives (blue point in Fig. 3d), because of the greater number of times the other asks for help. In general, a lower quality individual will ask for help more often, and there is also substantial random variation in how often individuals fail in foraging and need help. A random sample of relationships from the simulation illustrates that there is notable reciprocity in the relationships (grey points in Fig. 3d; the Spearman correlation between total donated and received is *r*_S_ = 0.77 for this simulation), but it is still common that a lack of strict reciprocation gives rise to a negative helping balance for one of the partners, causing a reduction in the amount it donates (cf. Fig. 1d), and this is the reason for the band-like pattern in Fig. 3d.

**Figure 3:**
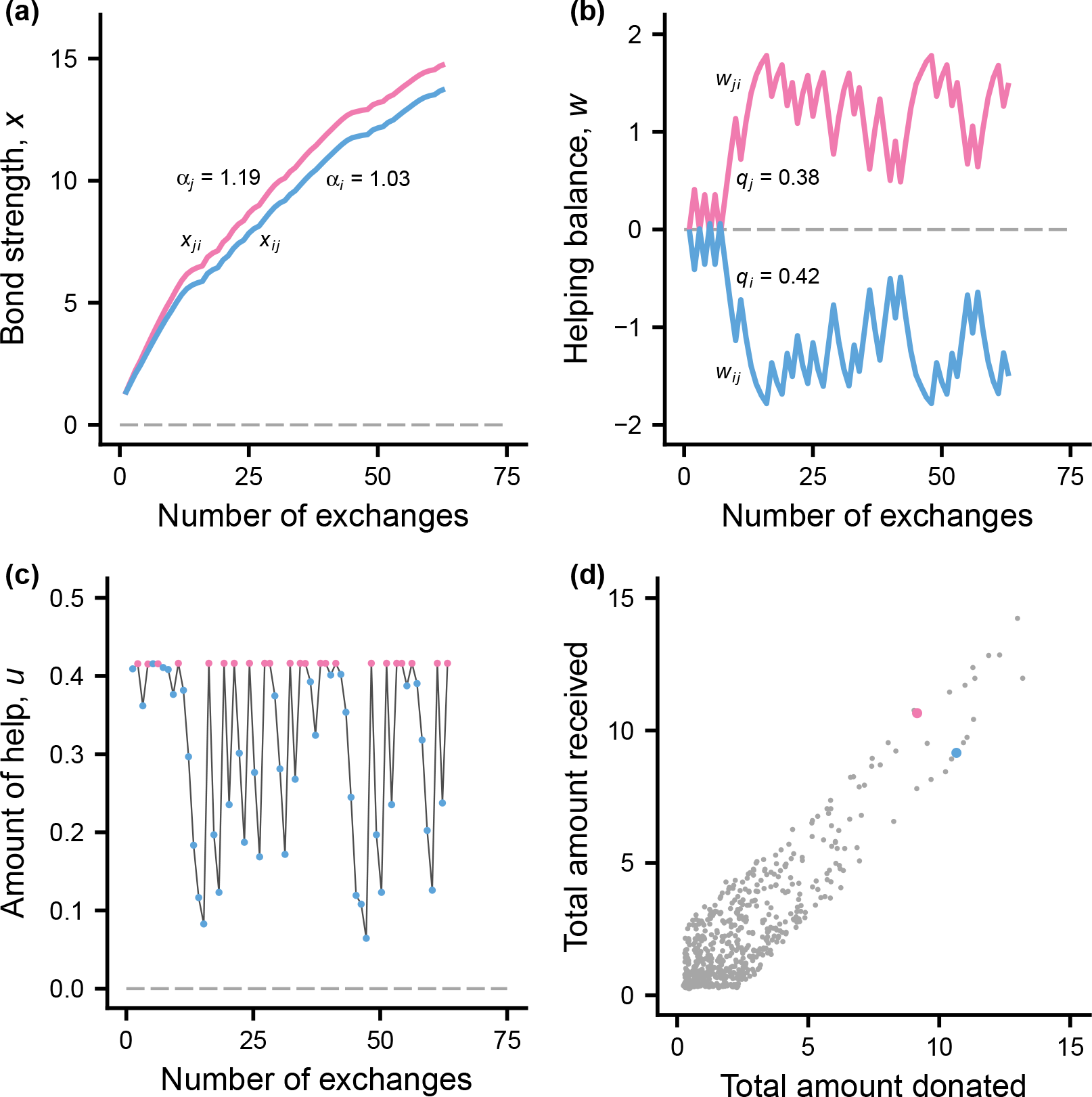
Example of bond dynamics and helping for a pair *i* and *j* with many helping exchanges. In the panels, blue and red indicate *i* and *j*. **(a)** The build-up of bond strength *x*_*ij*_ for *i* and *x*_*ji*_ for *j*. The genetically determined bond strength learning rates *α*_*i*_ and *α*_*j*_ appear near the corresponding curves. **(b)** The total helping balance *w*_*ij*_ for *i* and *w*_*ji*_ for *j*, together with the phenotypic quality values *q*_*i*_ and *q*_*j*_ for the example. **(c)** The amounts of help *u*_*ij*_ and *u*_*ji*_ donated by one individual to the other. **(d)** The colour coded points show the total amounts donated and received between *i* and *j*. The additional grey points show a sample of 1000 partnerships from those with helping at least once in each direction. The data come from the same simulation as in Fig. 2.

A striking aspect of the pattern of helping seen in Fig. 3c is that individuals donate close to their maximum amounts at the start of a relationship. The reason is that the bond strength increment (trait ∆*y*_new_, Fig. 1c) evolved to a large positive value (see Table S1 for evolutionary equilibrium trait values). As a consequence, individuals have a strong tendency to initiate relationships with new permanent group members, both in terms of requesting help and amounts donated, and do not follow a raise-the-stakes procedure (but see below for ‘visitor’ individuals).

Concerning the structuring into places, Fig. 4a shows the distribution of estimates of association when help is exchanged in pairs with previous helping in both directions (established partners). The distribution contains mostly high association values. When including all helping exchanges, lower association values are added to the distribution (Fig. 4b), making it similar to the association between randomly chosen pairs from the same place (Fig. 4c).

**Figure 4:**
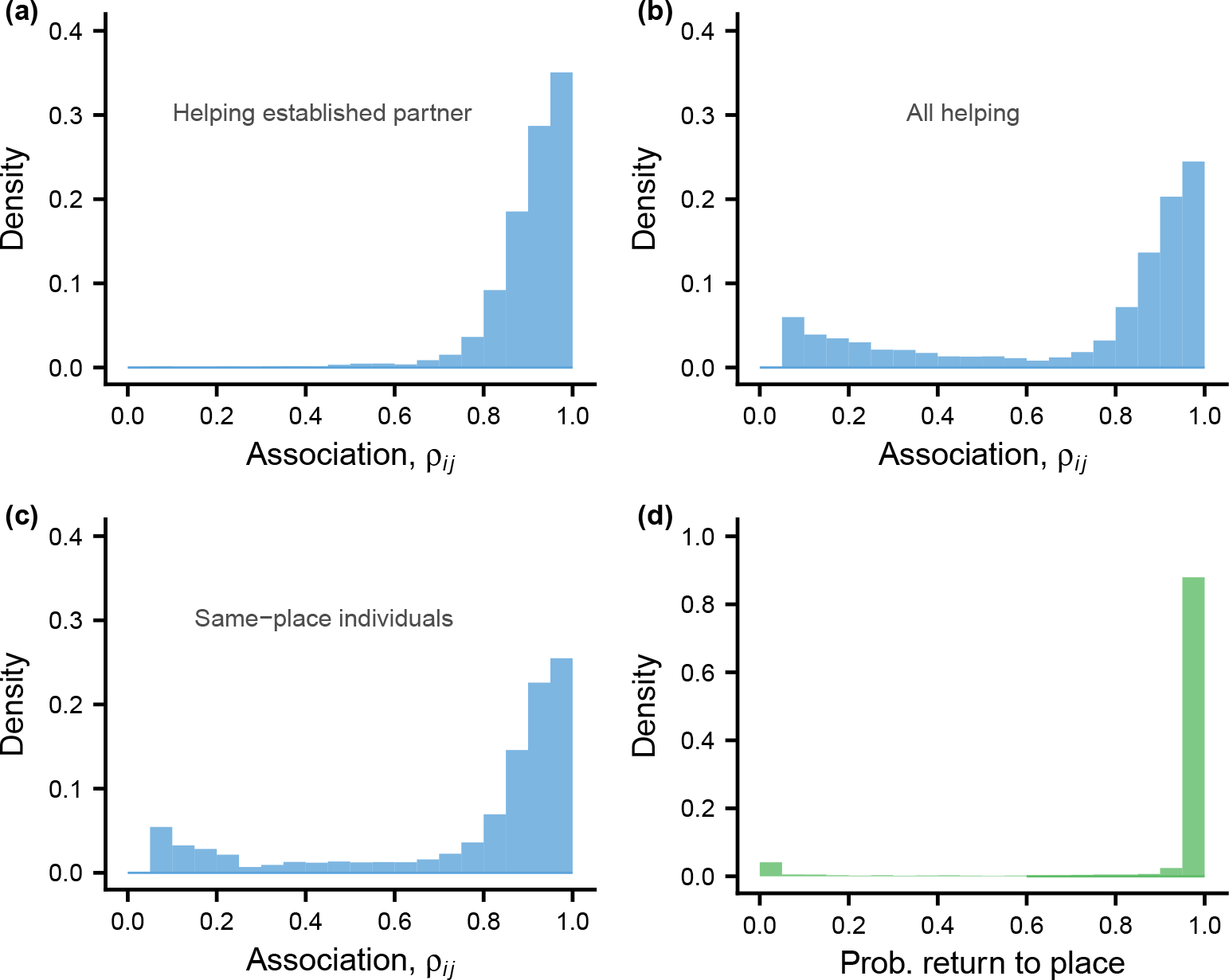
The distribution of estimates *ρ*_*ij*_ of associations between individuals, for different samples of pairs *i* and *j*, and the degree of place preference shown by individuals, measured as their probability to return to the current place in the next period. **(a)** Distribution of *ρ*_*ij*_ for helping between established partners, in the sense of recipient and donor having previously exchanged help in both directions. **(b)** Distribution for all helping interactions. **(c)** Distribution for individuals in the same place, regardless of helping interactions. **(d)** Distribution of the probability to return to the current place in the next period, shown as a density histogram. The data come from the same simulation as in Fig. 2.

A strong tendency to prefer places with higher average bond strength evolved in the simulation (Table S1). As a consequence, the probability that an individual returns next day to its current place is concentrated near 1.0 (Fig. 4d). The model implements a constraint on the choice of place, such that an individual ends up in a random place with a small probability (0.05). This is meant to represent variability in the particular ways that individuals distribute themselves over places on a given day, and allows individuals to encounter all group members. The constraint explains the small bump near zero in Fig. 4d; these individuals ended up in a random place and will mostly not return there the next day.

### The size of the social neighbourhood

A relatively small size of the social neighbourhood is needed for substantial helping to evolve in our model (Fig. 5). For larger group sizes *N*, the social neighbourhood can be a place with on average a small number *G* of individuals (Fig. 5a, b, c), or it can be a smaller group with a single place (Fig. 5d). For larger social neighbourhoods, the trait *h*_a*i*_, giving the maximum amount donated, evolves to a small value, resulting in little helping, as illustrated in Fig. 5f. Se also Figs. S2 and S3 for illustrations of the total amounts received and donated for the cases in Fig. 5.

**Figure 5:**
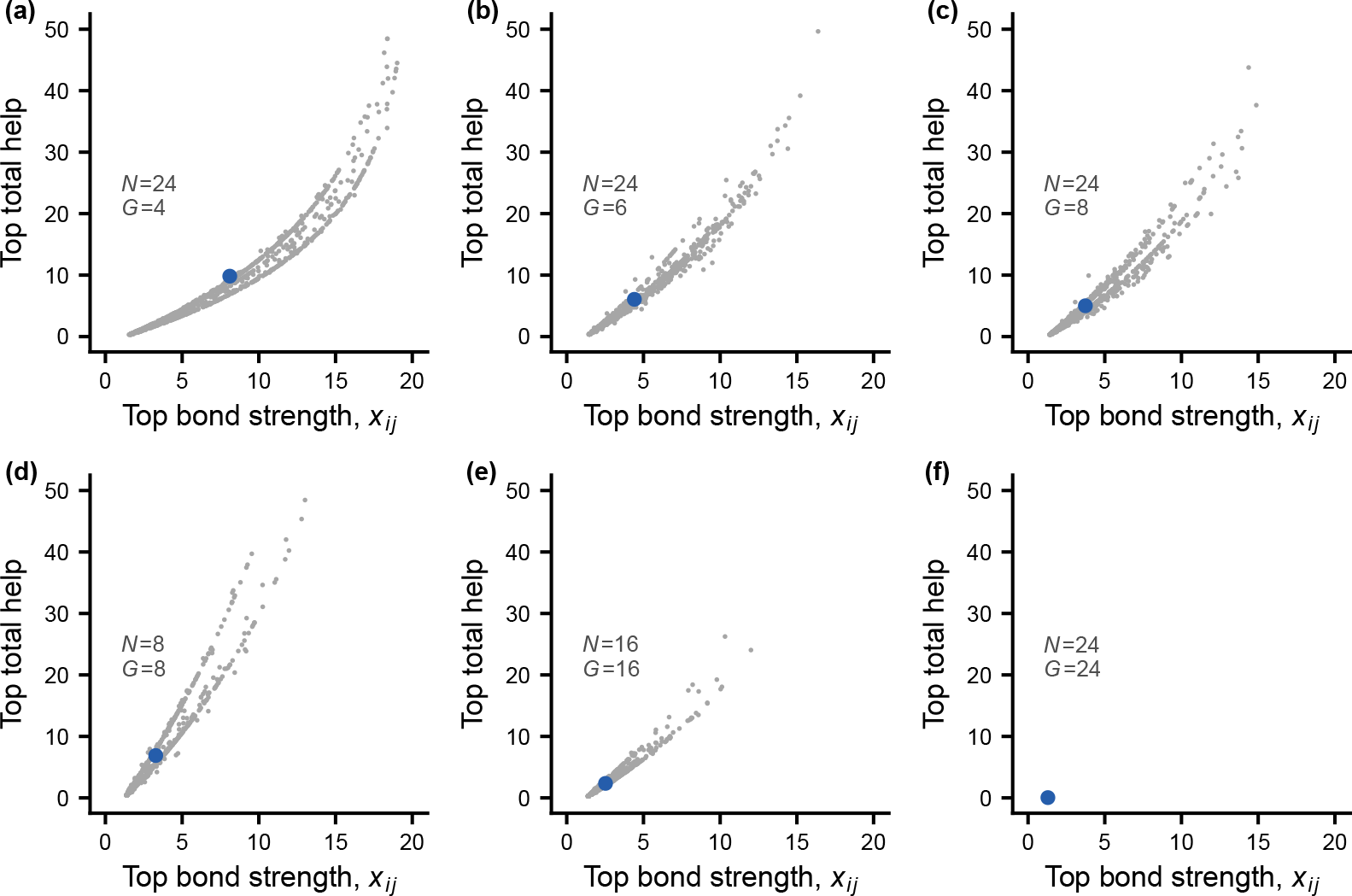
Effects of the size of the social neighbourhood on bond formation and helping. The top bond strength for an individual is the highest value of *x*_*ij*_ among the partnerships the individual has initiated over its lifetime, and the top total help is the sum of all help donated and received in this top partnership. The grey points in the panels show random samples (of size 1000) of individuals from evolutionary simulations, and the larger blue points show the sample means. Each simulation has a population size of at least 4000, split into groups of size *N*. Each group has one or more places where individuals can associate, and *G* is the expected number of individuals in a place. Panels **(a), (b)**, and **(c)** show cases with group size *N* = 24, split up into 6, 4, or 3 places (so that *G* = 4, 6, 8), and panels **(d), (e)**, and **(f)** show cases with different group sizes *N* = 8, 16, 24 and a single place (so that *G* = *N*). Note that helping evolved to near zero in panel **(f)**. Each simulation was run at evolutionary equilibrium for 4000 periods, and the random samples are of individuals born in the earlier part of this interval.

Part of the explanation for the reduction or elimination of helping with larger social neighbourhoods is that individuals evolve a strong tendency to initiate bonds with new group members (the trait ∆*y*_new_ evolves to a large positive value; Table S1). Individuals thereby gain the short-term advantage of a large network of potential partners, but this has the evolutionary consequence of reducing the amount of help donated. The explanation can be checked by fixing the trait ∆*y*_new_ at a negative value, causing individuals to prefer to request help from and donate help to established partners, and then see if helping evolves. The result is that notable helping with social bonds evolves for the group size in Fig. 5f, which is illustrated in Figs. S4 and S5 (see also Table S1). In this situation, there are high bond strength learning rates and relationships show an initial increase in helping amounts, corresponding to raising the stakes (Fig. S5).

To further investigate evolution of the tendency to initiate new bonds, we developed a variant of the model where individuals from one group sometimes visit another group, but return to their original group after a number of days (see Supplements for details). Evolutionary simulations for this model variant show that individuals avoid forming bonds between groups, but still evolve a strong tendency to initiate bonds with new within-group individuals (Table S2). As a consequence, essentially no help is exchanged between members of different groups (Fig. S6).

### Marginal costs and benefits

Helping amounts in our model are quantitative variables, so we can compute the increase or decrease in mortality from a slight change in the amount of help donated, for any particular situation (see Supplements). Performing this for the hypothetical case of exceedingly small amounts of help, we find a large marginal benefit-cost ratio of around 80. For the case illustrated in Fig. 3, with amounts around *u* = 0.4, the marginal benefit-cost ratio is around 8 for average quality individuals. These are high benefit-cost ratios, and are typical for our model of helping with social bonds.

### Survival interdependence, reciprocity, and partner choice

One possible explanation for the evolution of helping is that individuals depend on each other for help when help is needed. As a consequence, they would have an interest in promoting each other’s welfare. The interdependence would be strongest in small groups and become increasingly diluted for larger group sizes. Another explanation is that reciprocity promotes helping, so that individuals help in order to receive reciprocal help when they themselves need it. For our model, reciprocity is more important than survival interdependence in promoting the evolution of helping. For the smallest possible group size of *N* = 2, substantial helping evolves when reciprocity operates, but there is very little helping when reciprocity is prevented, by fixing the trait *h*_s*i*_ at zero (compare cases 9 and 10 in Table S3).

The concept of a social bond presupposes individual recognition, but one can still investigate what happens to the evolution of helping if an individual treats all group members as the same, as in generalized reciprocity [17]. Implementing this in the model (see Supplements for details), and performing evolutionary simulations of situations with group sizes and places as those Fig. 5, the maximum help trait *h*_a*i*_ evolved to near zero, except for the case with a small group size of *N* = 8 (Fig. S7 and Table S3). Thus, individual recognition is important for the evolution of helping with social bonds. Comparing Fig. S3 with Fig. S7, an interpretation is that partner choice, by way of the social bond mechanism, contributes to the evolution of helping. Thus, partner choice and reciprocity appear to be the two main mechanisms promoting the evolution of helping in our model.

## Discussion

Behavioural mechanisms are the main ingredients in our model of helping with social bonds, and we study the evolution of traits that influence these mechanisms through individual-based simulation. An important mechanism is the dynamics of social bonds, which we describe as a psychological process qualitatively similar to associative learning, with a learning-rate trait *α*. At present, the psychology of social bonding is not well enough developed to settle if the analogy to learning is exact, or if bond formation is a similar but separate psychological mechanism. The social bond dynamics in our model has an added element that determines how potential new partners are handled. This element (the trait ∆*y*_new_) influences the growth and eventual size of an individual’s social network. We found that there is selection for individuals to incorporate new (permanent) group members into their social network (∆*y*_new_ evolved to positive values; Fig. 1, Table S1). On the other hand, we found that there is selection for individuals to choose to spend time in subgroups (places) where they are likely to find their socially bonded partners and can maintain the association to these partners (the place choosiness trait *β* evolved to positive values; Figs. 1, 4, Table S1). Our analysis thus identifies a tension between, on the one hand, gaining security by maintaining and strengthening existing bonds and, on the other hand, gaining security by enlarging the social network.

This tension is responsible for a limitation on the evolution of helping with social bonds that we identified. We found that small social neighbourhoods are required (Fig. 5), because for large social neighbourhoods the maximum level of helping (the trait *h*_a_) evolves to a small value. In situations with small social neighbourhoods, where helping through social bond formation does evolve, we found that a combination of reciprocity (by way of the trait *h*_s_) and partner choice (based on the strength of social bonds) is what maintains helping.

Our model is inspired by observations on female vampire bats [4, 18, 19, 20], specifically our assumptions that food donations go from successful to unsuccessful foragers (with a fairly high chance of being successful), that individuals in need can request help more than once, that groups are structured into one or more resting places (day roosts), and that the survival benefits and costs of food donations have approximately exponential shapes (Fig. 2a). For simplicity, we did not model the sex differences observed in vampire bats, instead assuming that all individuals are equally involved in food sharing. We also did not model the natural within-group recruitment of female offspring, instead implementing hypothetical offspring recruitment from the entire population, in order to prevent evolutionary effects of relatedness from contributing to our results.

We might then compare the qualitative results from our model with field and experimental observations on female vampire bats. First, for estimates of same-place associations, there is the similarity that food-sharing pairs tend to have higher association (compare Fig. 4 with [4, 21, 22]). Second, our analysis revealed a form of reciprocity that is neither immediate nor very strict (Fig. 3), and this also seems to hold for reciprocity in female vampire bats [5]. Third, our finding of an evolved tendency for individuals to incorporate new permanent group members into their social network is in line with experiments showing that non-kin food sharing by female bats expands their social network, which reduces their future risk of not receiving help when they need it [23, 24], and this might explain the evolution of non-kin food sharing [25]. Finally, we did not model the fusion of different groups, but the results from our model version with short-term visitors between groups shows some similarity to the experimental fusion of two groups of bats [14] (currently it is not known if fusions occur in the wild). Specifically, we found that group members avoid forming bonds with visitors (Fig. S6, Table S2), and we also analysed a special case of a negative ∆*y*_new_ (Fig. S5, Table S1),

While our current results (Fig. 3c) do not support the idea of raising-the-stakes cooperation [13, 14], different assumptions about how new individuals are recruited into groups might change this. Raising-the-stakes could be interpreted as a form of defence against potential exploiters that request help and then move on. As there will be costs associated with delaying the exchange of help, there needs to be a noticeable presence of such exploiters for a strategy of initial caution to be evolutionarily stable (see [26] for a discussion of this point).

There is one previous evolutionary model investigating how individuals might form social bonds when deciding on the distribution of help to other group members [15]. The work uses the idea of a Bayesian update (of the probability of donating help when requested) as inspiration for a social bond mechanism. There is some similarity with our approach to social bond dynamics, in that both models describe the build-up of bond strength, but there are also major differences, for instance in how the helping of individuals in need is implemented (e.g., from our analysis, new permanent group members receive substantial help, Fig. 3c, which is not the case for the analysis in [15]). At present, observations that could shed light on the lifetime dynamics of helping in vampire bats seem not to be available. Another major difference is that the maximum amount of help is an evolving trait in our model, underpinning our conclusion that small social neighbourhoods are required for substantial helping to evolve.

More generally, social bonds are found in several groups of mammals and some birds [2, 3, 27, 28]. Primate social bonds are the most studied, and an overall conclusion is that bonded (socially integrated) individuals have advantages in terms of health, survival, and reproduction [2, 3]. The acts of helping tend to be low-cost behaviours [29], while the advantages of receiving help can be substantial, and this also holds for our results.

The study of the evolution of cooperation is a very large field, in which a range of ideas continue to be evaluated [12]. The importance of reciprocity for helping in social groups is one of the most discussed of these ideas, first dealt with by Trivers more than 50 years ago [7]. While immediate and strict reciprocity appears to be rare [30], there is still the possibility that reciprocity operates between group members over a longer time frame [29, 11], and this is broadly in agreement with our results. There are several other important possibilities that we did not include in our analysis, for instance relatedness and indirect benefits [12], the punishment of defectors [31], and the abandonment of less profitable interactions [32]. The focus of our model is on the particular issue of how helping with social bonds might operate and evolve.

In contrast to traditional analyses of cooperation, such as those based on the iterated Prisoner’s Dilemma, our approach is to incorporate specific and potentially realistic cognitive and behavioural mechanisms into game-theory models. This approach is a new development of game theory in biology [33, 1]. It introduces a certain complexity of assumptions about traits, mechanisms, and life histories, but it has the crucial advantage of bringing modelling and observation into closer contact. In our view, an energetic pursuit of such contact is essential for progress in the challenging study of social interactions, including the evolution of cooperation in social groups. If we are to understand complex phenomena such as social bonds from an evolutionary point of view, we need a game theory that examines the mechanistic underpinnings of decision-making.

## Methods

An individual *i* has five genetically determined traits: *α*_*i*_ (bond strength learning rate), *β*_*i*_ (bond strength place sensitivity), ∆*y*_new *i*_ (bond strength increment for new partner), *h*_a*i*_ (asymptotic helping amount), and *h*_s*i*_ (helping balance sensitivity). Evolutionary equilibrium values of these traits for different simulations appear in Supplementary Tables S1, S2, and S3, together with a full and detailed model description, and with notation for the model in Table S4. In order to allow mutations to efficiently explore the multidimensional trait space, individuals are assumed to be haploid, reproducing sexually such that two parents form a diploid that then produces a haploid offspring, with mutation and recombination. Mutational distributions have long-tailed, Laplacian distributions (see Supplements).

For an individual in need (foraging success *z*_*i*_ = 0), if there is more than one other individual available and estimated to have succeeded in foraging (*z*_*j*_ = 1), the individual chooses to ask help from *j* with a probability proportional to exp(*y*_*ij*_), where *y*_*ij*_ is the effective bond strength to *j*. For a choice between two individuals, with a bond strength difference of ∆*y* between them, the probability of choice then becomes

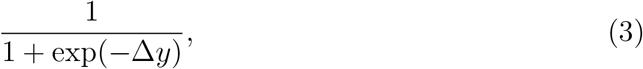

which is illustrated in Fig. 1b. Choices between places are similarly influenced by an individual’s estimate of the average effective bond strength to other individuals encountered in different places. There is a small probability *ϵ*_p_ that the individual ends up in a random place (we used *ϵ*_p_ = 0.05), and otherwise the probability to choose place *k* is proportional to exp(*β*_*i*_*ŷ*_*ik*_), where *ŷ* _*ik*_ is the estimate. For two places to choose between, and with ∆*y* the difference between them in estimated average bond strength, the probability of choice becomes

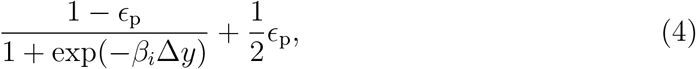

which is illustrated as a dashed line in Fig. 1b (with *β*_*i*_ = 7.9; see Supplements for further details, including a density dependence of the choice).

The amount of help individual *i* donates when *j* request help depends on several factors, and is expressed as a product *u*_*ij*_ = *H*_*y*_*H*_*w*_. The dependence on the effective bond strength is

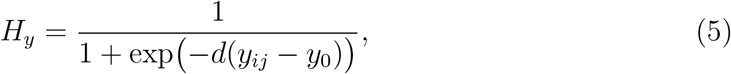

where *d* and *y*_0_ are parameters (we used *d* = 8.0 and *y*_0_ = 1.5). This is illustrated in Fig. 1c, including the effect ∆*y*_new *i*_ of donating for the first time to new individual. There is also a dependence on the helping balance *w*_*ij*_ with *j*, as well as on the quality *q*_*i*_, the foraging success state *z*_*i*_, and the current resource balance *ζ*_*i*_ of *i*, given by

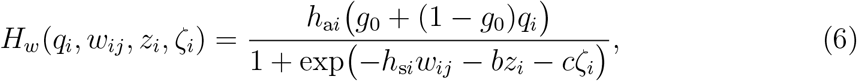

where *h*_a*i*_ and *h*_s*i*_ are genetically determined traits and *g*_0_, *b* and *c* are parameters (*b* and *c* also appear in the mortality function below). The function *H*_*w*_ is illustrated in Fig. 1d. Finally, the day-to-day mortality rate for an individual *i* is

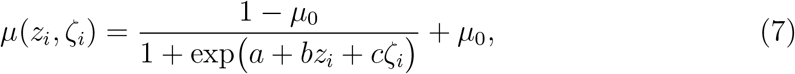

where *a, b, c* and *µ*_0_ are parameters (we used *a* = 2, *b* = 4.5, *c* = 3.0, and *µ*_0_ = 0.002). Donating or receiving help changes the resource balance *ζ*_*i*_ (e.g., receiving the amount *u* increases *ζ*_*i*_), and thus the mortality rate, and this is illustrated in Fig. 2a.

## Acknowledgements

We thank Peter Hammerstein for helpful comments.

## Supplementary information

### Supplementary figures

**Figure S1:**
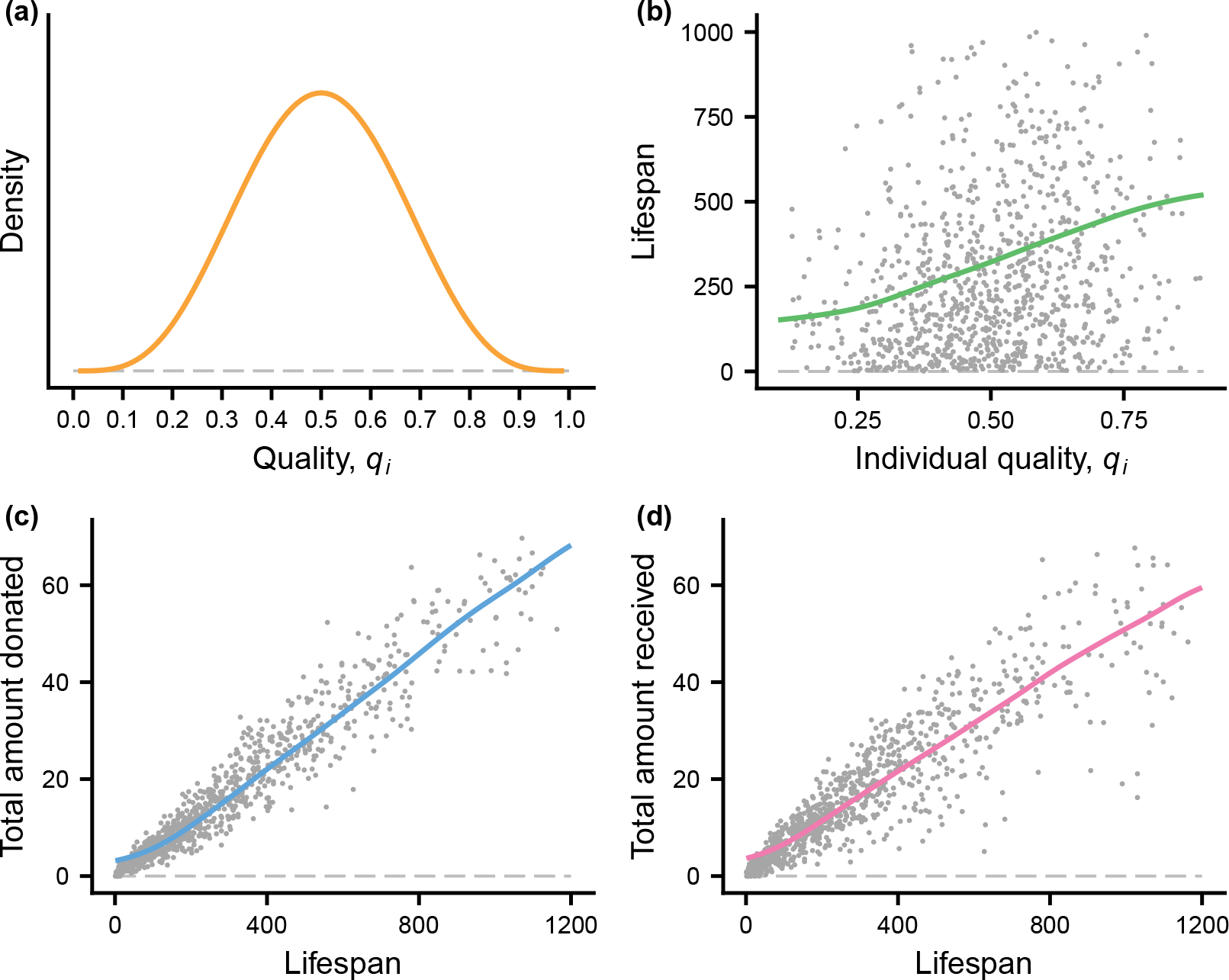
**(a)** The distribution of individual quality (*q*_*i*_) is given by a beta distribution with shape parameters equal to 5, B(5, 5). **(b)** Lifespan vs. individual quality for a random sample of 1000 individuals from those analysed in Fig. 2, together with the fitted curve in Fig. 2b. **(c)** and **(d)** Total amounts of help donated and received vs. lifespan for the sample, together with the fitted curves from Fig. 2c.

**Figure S2:**
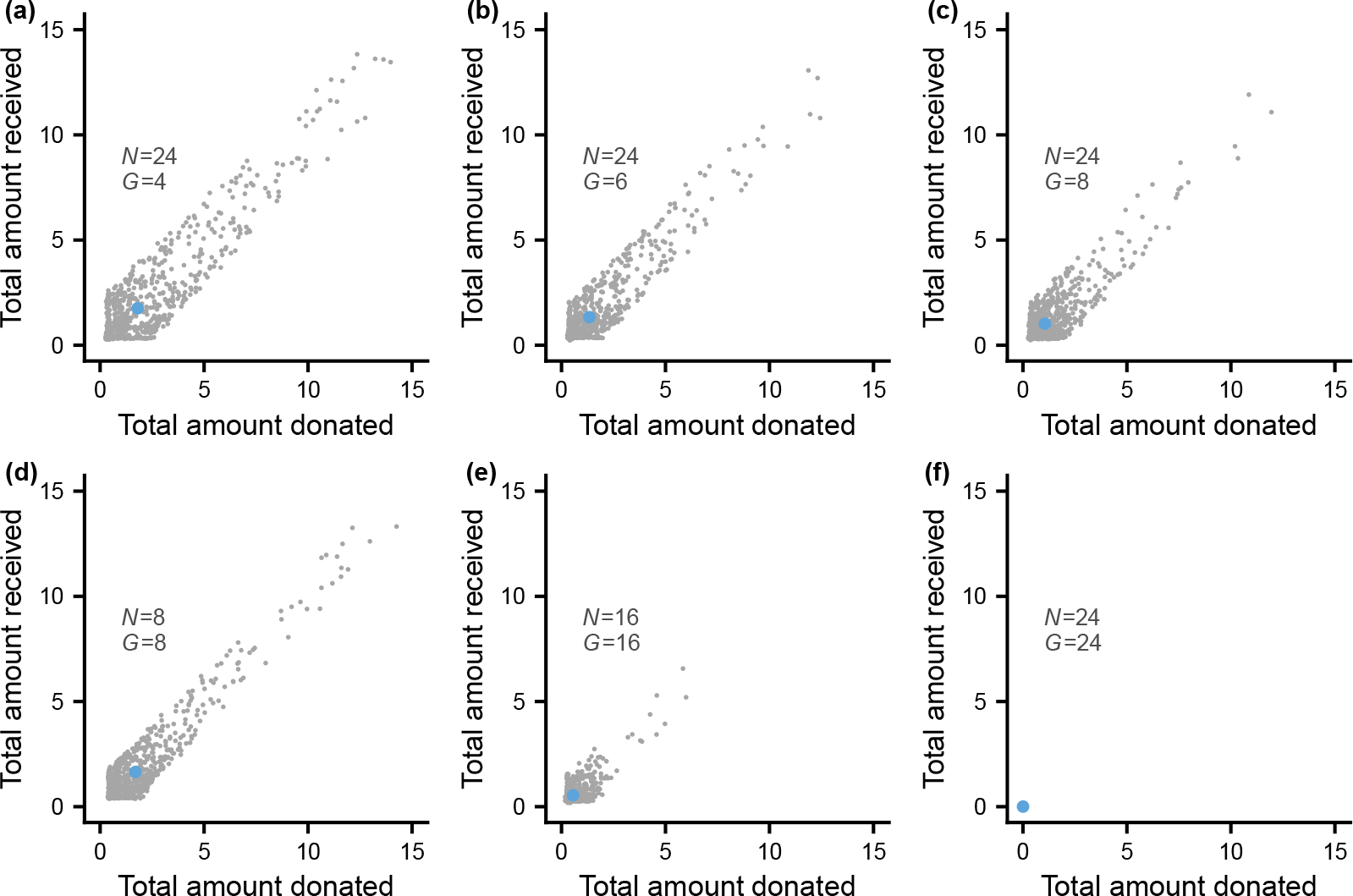
Effect of the size of the social neighbourhood on the total amount of help in relationships with at least one exchange in each direction. The grey points in the panels show total received vs. total donated in random samples (of size 1000) of relationships from evolutionary simulations. The larger blue points show the sample means. Each simulation (used in Fig. 5) has a population size of at least 4000, split into groups of size *N*. Each group has one or more places where individuals can associate, and *G* is the expected number of individuals in a place. Panels **(a), (b)**, and **(c)** show cases with group size *N* = 24, split up into 6, 4, or 3 places (so that *G* = 4, 6, 8), and panels **(d), (e)**, and **(f)** show cases with different group sizes *N* = 8, 16, 24 and a single place (so that *G* = *N*). Note that helping evolved to near zero in panel **(f)**. Each simulation was run at evolutionary equilibrium for 4000 periods, and the random samples are of individuals born in the earlier part of this interval (thus including those with long lifespans).

**Figure S3:**
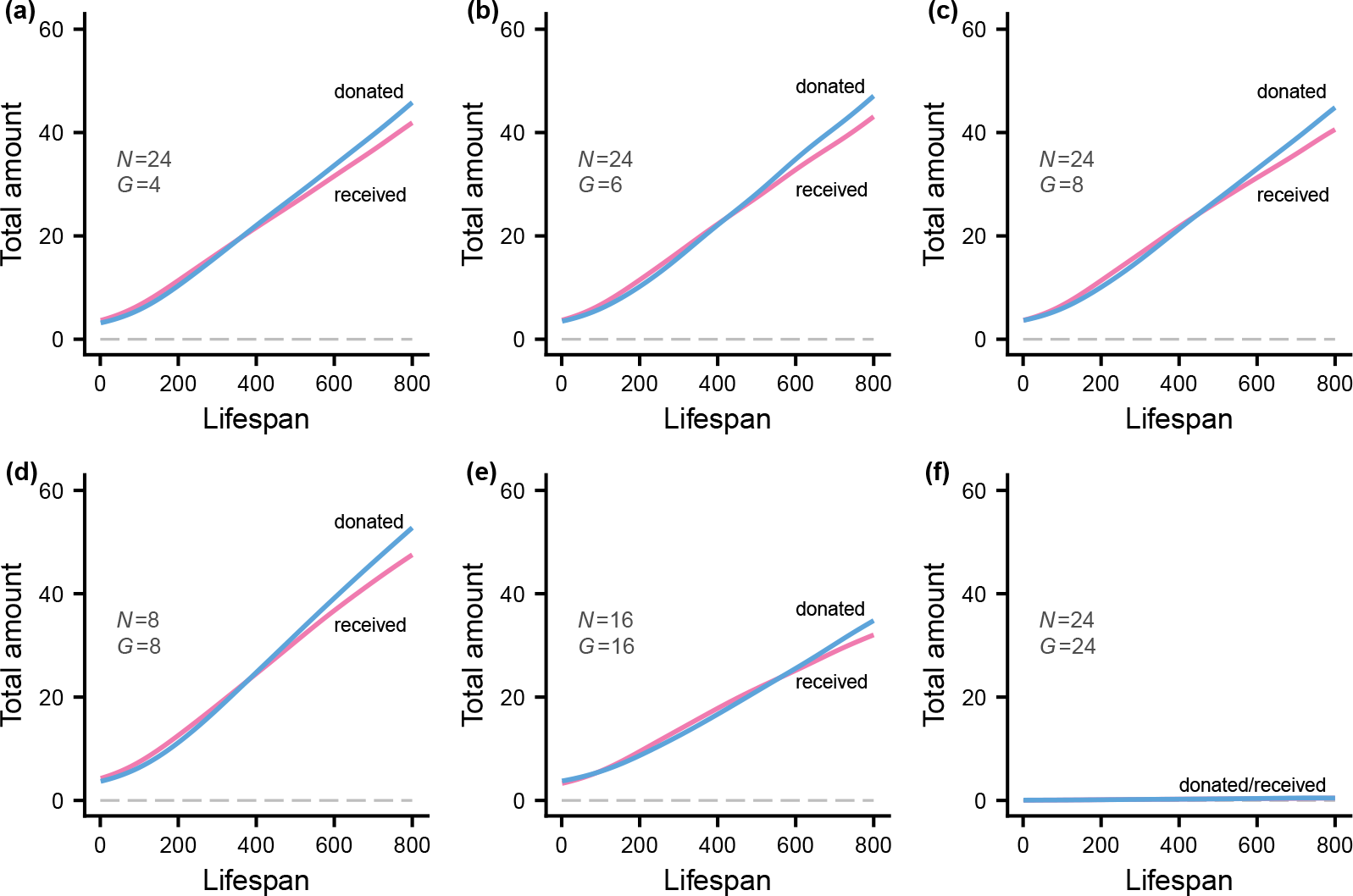
Effect of the size of the social neighbourhood on the expected total amount of help donated and received (to and from all other individuals) as a function of an individual’s lifespan. Each simulation (used in Fig. 5) has a population size of at least 4000, split into groups of size *N*. Each group has one or more places where individuals can associate, and *G* is the expected number of individuals in a place. Panels **(a), (b)**, and **(c)** show cases with group size *N* = 24, split up into 6, 4, or 3 places (so that *G* = 4, 6, 8), and panels **(d), (e)**, and **(f)** show cases with different group sizes *N* = 8, 16, 24 and a single place (so that *G* = *N*). Note that helping evolved to near zero in panel **(f)**. Each simulation was run at evolutionary equilibrium for 4000 periods, and the random samples are of individuals born in the earlier part of this interval (thus including those with long lifespans). The curves are kernel smoothing fits and are based on ca 12 000 individuals.

**Figure S4:**
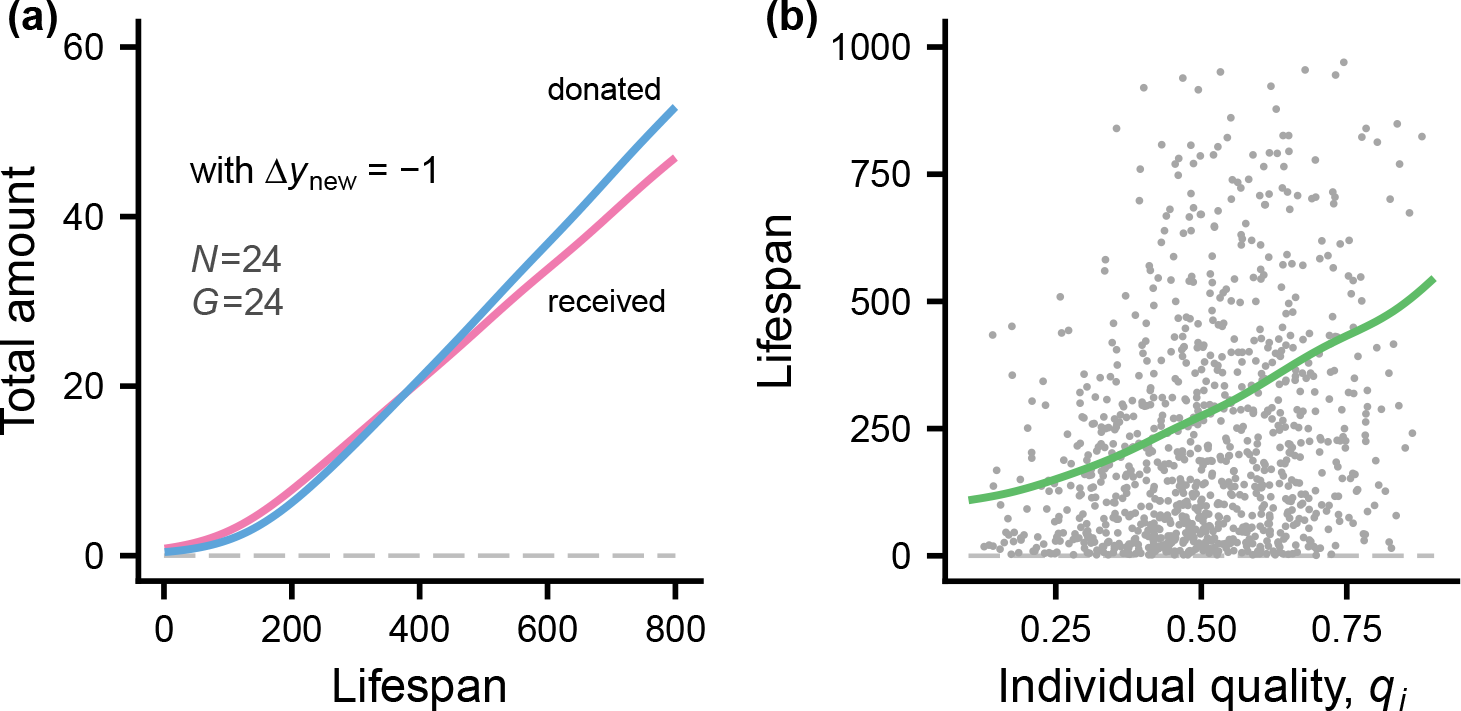
Evolution of helping when there is a tendency to avoid forming bonds with new individuals. This is achieved by simulating evolution with the trait ∆*y*_new *i*_ fixed at *−*1. **(a)** Total amounts of help donated and received as a function of lifespan. **(b)** Lifespan vs. individual quality for a random sample of 1000 individuals, together with a fitted curve. The simulation has a population size of 4200, split into groups of size *N* = 24 with a single place (*G* = 24). The curves are kernel smoothing fits and are based on ca 12 000 individuals, including those with long lifespans.

**Figure S5:**
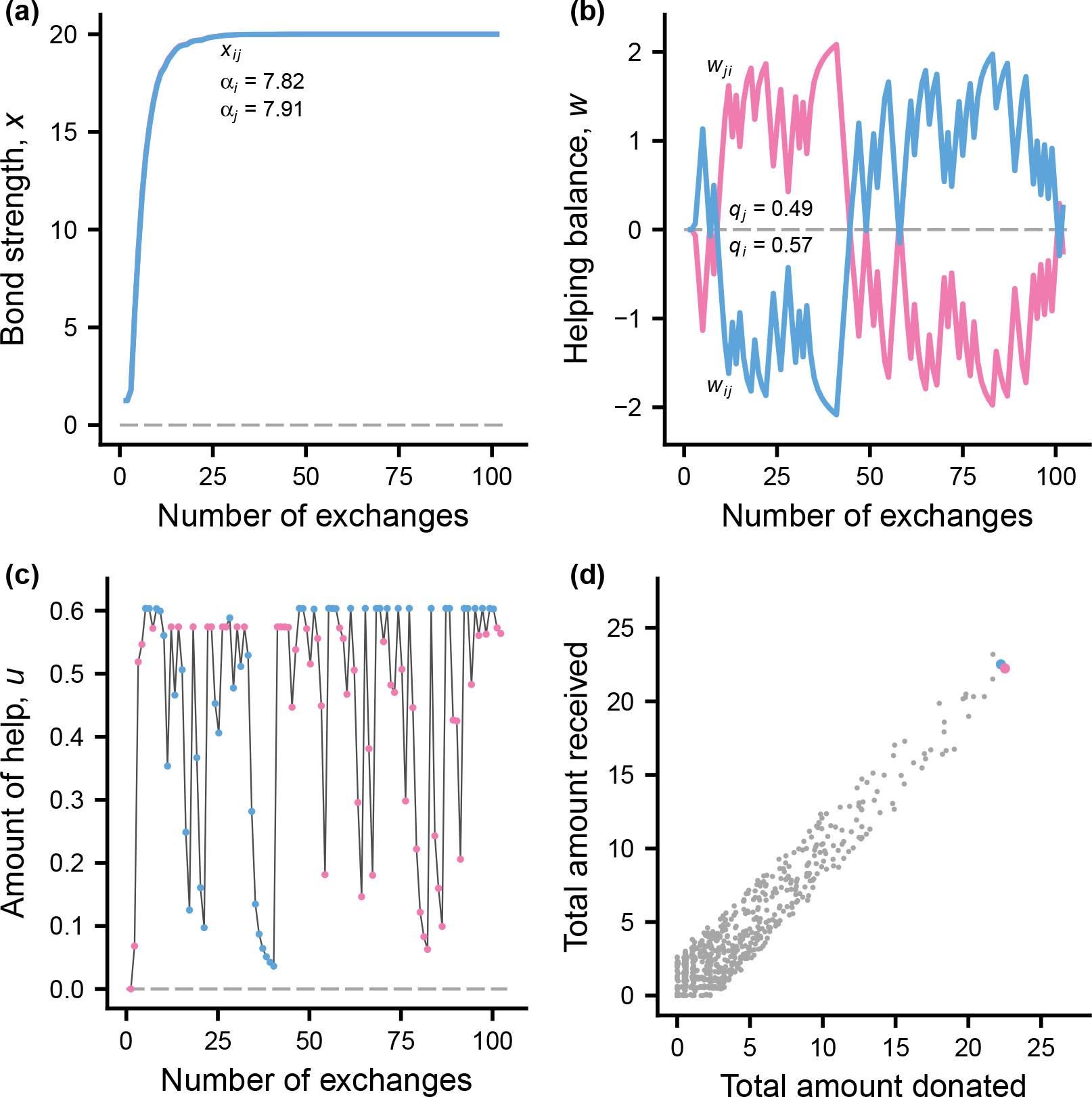
Example of bond dynamics and helping for a pair *i* and *j* with many helping exchanges, from the simulation with the trait ∆*y*_new *i*_ fixed at *−*1 (see Fig. S4). In the panels, blue and red indicate *i* and *j*. **(a)** The build-up of bond strength *x*_*ij*_ for *i* (which overlaps with *x*_*ji*_ for *j*). The genetically determined bond strength learning rates *α*_*i*_ and *α*_*j*_ are shown in the panel. **(b)** The total helping balance *w*_*ij*_ for *i* and *w*_*ji*_ for *j*, together with the phenotypic quality values *q*_*i*_ and *q*_*j*_ for the example. **(c)** The amounts of help *u*_*ij*_ and *u*_*ji*_ donated by one individual to the other. **(d)** The colour coded points show the total amounts donated and received between *i* and *j*. The additional grey points show a sample of 1000 partnerships from those with helping at least once in each direction. The data come from the simulation shown as case 7 in Table S1.

**Figure S6:**
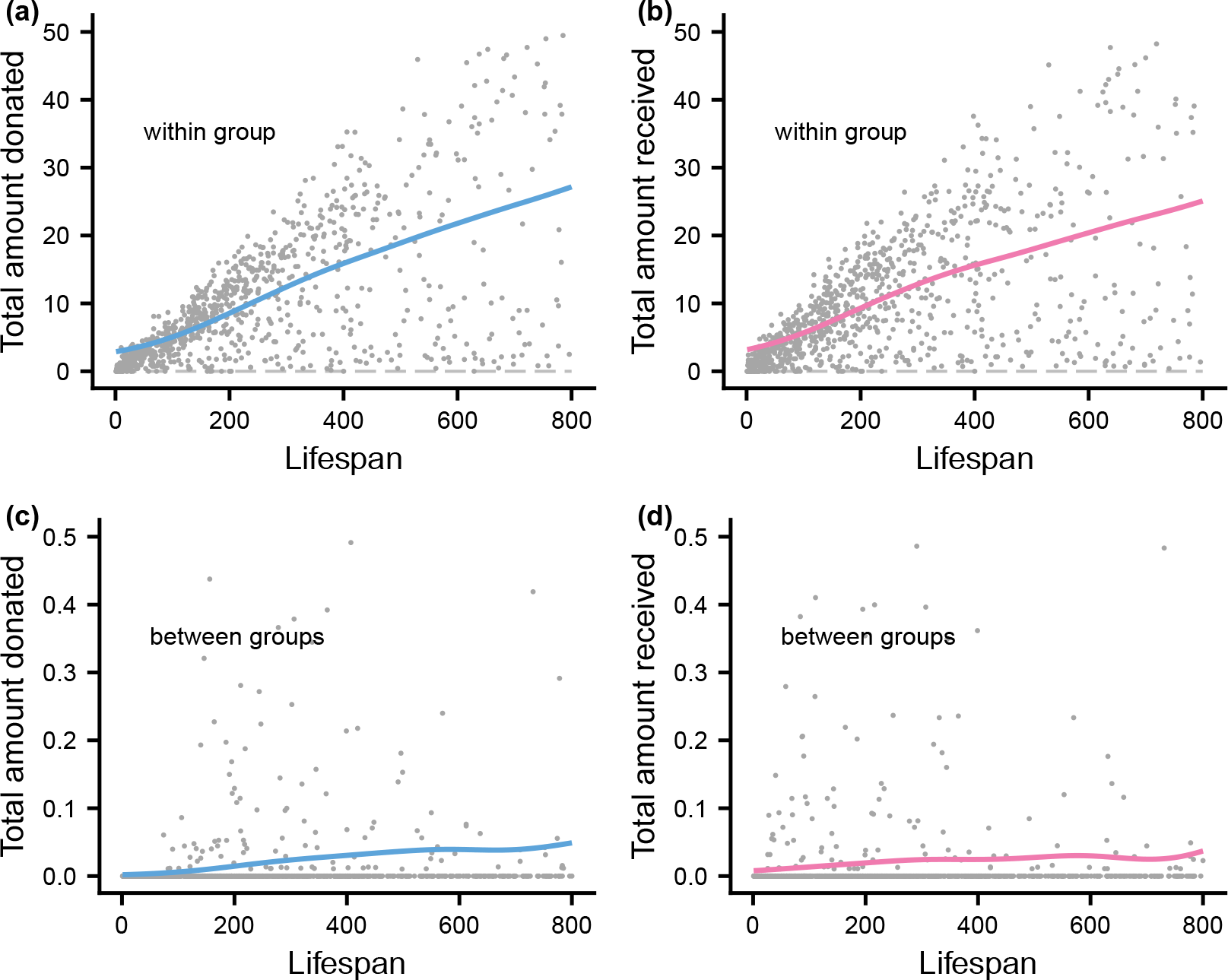
Total amounts donated and received as a function of lifespan for the model variant where some groups exchange visitors over a short period (20 days). The visitors subsequently return to their original group. **(a)** and **(b)** Total amounts of help donated and received between members of the same original group as a function of lifespan. **(c)** and **(d)** Total amounts of help donated and received between members of different original groups as a function of lifespan. Note the difference in scale on the *y*-axes.

**Figure S7:**
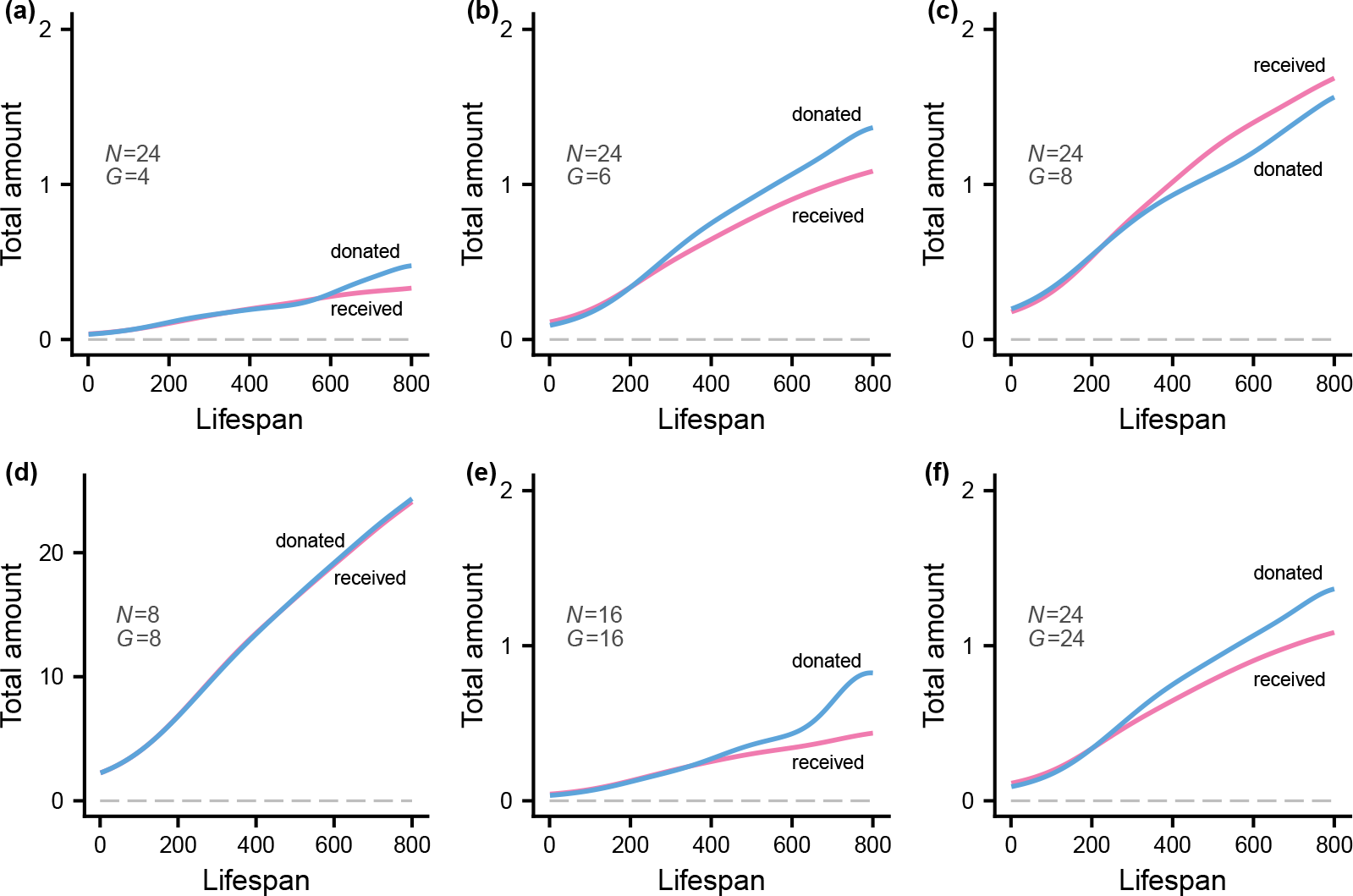
Effect of the size of the social neighbourhood, as in panels **(a), (b), (c)** of Fig. S3, but for evolutionary simulations without individual recognition. Note that the scale on the *y*-axis differs from that in Fig. S3. With individual recognition (Fig. S3), the amounts of help exchanged are much larger than here. Each simulation has a population size of 4200, split into groups of size *N* = 24, with one or more places, and *G* is the expected number of individuals in a place. Removing individual recognition means that an individual treats all other individuals as the same, having a social bond to this collective, and there is no basis for preferring one place to another. The curves are kernel smoothing fits and are based on ca 12 000 individuals. Note the different scale on the *y*-axes in panel **(d)**.

### Supplementary tables

**Table S1:**
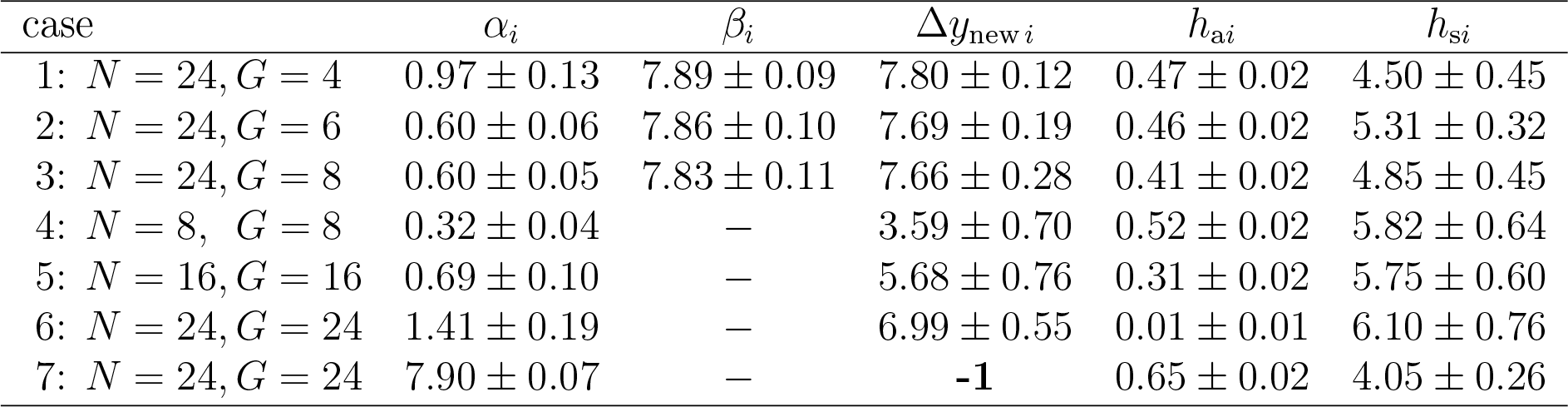
Trait values (mean *±* SD over 100 simulations, each over 100 000 days) for 7 different cases of individual-based evolutionary simulations of helping with social bonds. The parameters that vary between cases are the group size (*N*) and the average number of individuals per place (*G*). For case 7, one trait was kept fixed.

**Table S2:** Trait values for a simulation with the model variant where groups sometimes exchange temporary visitors, with *N* = 16 and *G* = 8.

| case | $\alpha_i$ | $\beta_i$ | $\Delta y_{\text{new } i}$ | $\Delta y_{\text{vis } i}$ | $h_{ai}$ | $h_{si}$ |
| --- | --- | --- | --- | --- | --- | --- |
| 8: | $0.61 \pm 0.06$ | $7.29 \pm 0.54$ | $7.52 \pm 0.31$ | $-3.66 \pm 0.27$ | $0.46 \pm 0.01$ | $7.01 \pm 0.43$ |

**Table S3:** Trait values for 8 cases of individual-based evolutionary simulations of helping without individual recognition. The group size for cases 9 and 10 is *N* = 2 (*h*_s*i*_ is fixed at 0 for case 10), and the other correspond to cases 1 to 6 in Table S1.

| case | $\alpha_i$ | $\beta_i$ | $\Delta y_{\text{new } i}$ | $h_{ai}$ | $h_{si}$ |
| --- | --- | --- | --- | --- | --- |
| 9: $N = 2, G = 2$ | — | — | — | $0.80 \pm 0.04$ | $2.85 \pm 0.47$ |
| 10: $N = 2, G = 2$ | — | — | — | $0.02 \pm 0.02$ | **0** |
| 11: $N = 24, G = 4$ | — | — | — | $0.02 \pm 0.01$ | $4.42 \pm 0.76$ |
| 12: $N = 24, G = 6$ | — | — | — | $0.02 \pm 0.01$ | $3.79 \pm 0.65$ |
| 13: $N = 24, G = 8$ | — | — | — | $0.02 \pm 0.01$ | $4.67 \pm 0.57$ |
| 14: $N = 8, G = 8$ | — | — | — | $0.27 \pm 0.05$ | $3.52 \pm 0.33$ |
| 15: $N = 16, G = 16$ | — | — | — | $0.01 \pm 0.01$ | $1.90 \pm 0.35$ |
| 16: $N = 24, G = 24$ | — | — | — | $0.02 \pm 0.02$ | $4.56 \pm 0.79$ |

**Table S4:** Definitions and notation for the model.

| notation | definition or explanation |
| --- | --- |
| $N$ | size of local group |
| $K$ | number of places available to group members |
| $G$ | expected number individuals per place; $G = N/K$ |
| $z_i$ | current foraging success state (0/1) for individual $i$ |
| $q_i$ | phenotypic quality ($0 < q_i < 1$) for $i$; see Fig. S1a |
| $\zeta_i$ | current resource balance for $i$; $\zeta_i = 0.5q_i$ at start of day |
| $p_{si}$ | probability of foraging success; $p_{si} = p_{s0} + (1 - p_{s0})q_i$ |
| $p_{s0}$ | parameter; $p_{s0} = 0.8$ in simulations |
| $p_d$ | probability of detecting another's state $z_j$; $p_d = 0.98$ |
| $x_{ij}$ | bond strength between individuals $i$ and $j$ |
| $x_s$ | starting value of $x_{ij}$: $x_s = 1.25$ |
| $x_a$ | asymptotic (maximum) value of $x_{ij}$: $x_a = 20.0$ |
| $\rho_{ij}$ | same-place association between $i$ and $j$ |
| $y_{ij} = \rho_{ij}x_{ij}$ | effective (association adjusted) bond strength |
| $\hat{y}_{ik}$ | average effective bond strength for $i$ to place $k$ |
| $\alpha_i$ | bond strength learning rate for individual $i$ |
| $\beta_i$ | sensitivity to among place bond strength differences |
| $\Delta y_{\text{new } i}$ | bond strength increment for $i$ for a new partner |
| $h_{ai}$ | asymptotic helping amount for $i$ |
| $h_{si}$ | helping balance sensitivity for $i$ |
| $\Delta y_{\text{vis } i}$ | between-group version of $\Delta y_{\text{new } i}$ for model with visitors |
| $\mu(z_i, \zeta_i)$ | rate (per day) of mortality as a function of $z_i$ and $\zeta_i$ |
| $a, b, c, \mu_0$ | parameters for $\mu$; $a = 2.0$, $b = 4.5$, $c = 3.0$, $\mu_0 = 0.002$ |
| $u_{ij} = H_y H_w$ | amount of help donated by $i$ to $j$ |
| $H_y(y_{ij})$ | help as function of $y_{ij}$ |
| $d, y_0$ | parameters for $H_y$; $d = 8.0$, $y_0 = 1.5$ |
| $w_{ij}$ | helping balance: received minus donated between $i$ and $j$ |
| $H_w(q_i, w_{ij}, z_i, \zeta_i)$ | help as function of $q_i$, $w_{ij}$, $z_i$, and $\zeta_i$ |
| $g_0, b, c$ | parameters for $H_w$; $g_0 = 0.75$, $b = 4.5$, $c = 3.0$ |
| $\epsilon_p$ | probability of ending up in a random place; $\epsilon_p = 0.05$ |
| $\alpha_p$ | rate of updating of estimates $\rho_{ij}$, $\hat{y}_{ik}$, and $\hat{n}_{ik}$; $\alpha_p = 0.1$ |
| $\beta_d$ | sensitivity to density estimate in choice of place; $\beta_d = 4.0$ |
| $\hat{n}_{ik}$ | density estimate by $i$ for place $k$ |

### Model details

In order to achieve a self-contained presentation, we explain all model aspects here, while referring to the presentation in the main text.

Individuals spend their lives in groups of size *N*, with a substructure of *K* places available, so that on average there are *G* = *N/K* individuals per place. Time is divided into steps, where a time step might be one day. At the start of a day, individuals choose which place to go to, and the resource state *z*_*i*_ of each individual *i* is *z*_*i*_ = 1 with probability *p*_s*i*_ (the individual succeeded in foraging) and *z*_*i*_ = 0 with probability 1*−p*_s*i*_. An individual’s *p*_s*i*_ depends on its quality (see equation (S1) below). Only individuals *i* with *z*_*i*_ = 0 ask for help on a given day. They only ask help from individuals *j* they perceive have a resource state of *z*_*j*_ = 1. Individuals have a probability *p*_d_ of correctly detecting another’s resource state.

### Bond strength notation

The model combines two aspects of a social bond. One is the the accumulated history of helping between individuals *i* and *j*, which is denoted *x*_*ij*_. The other represents the recent history of same-place association between *i* and *j*, which is denoted *ρ*_*ij*_. The effective (association adjusted) bond strength *y*_*ij*_, influencing who to ask help from and how much help to donate, is the product of these: *y*_*ij*_ = *ρ*_*ij*_*x*_*ij*_. The qualitative effect of this formulation is that if one of two bonded individuals changes place, causing a decrease in *ρ*_*ij*_, if they again meet they will have a lower effective bond strength.

### Genetically determined traits

An individual *i* has five genetically determined traits: (1) a bond strength learning rate *α*_*i*_ used to update *x*_*ij*_, (2) a parameter *β*_*i*_ giving the degree of choosiness between places with different estimated bond strengths, (3) an increment ∆*y*_new *i*_ to the effective bond strength used when asking help from and donating to a new individual, (4) a helping asymptote *h*_a*i*_ and, (5) a helping sensitivity *h*_s*i*_ to the current helping balance.

### Individual quality

Individuals vary in phenotypic quality *q*_*i*_, which influences their probability *p*_s*i*_ of success in foraging. For individual *i*,

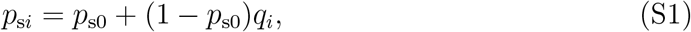

with, for instance, *p*_s0_ = 0.8. The quality *q*_*i*_ has a Beta distribution; we used a B(5, 5) distribution (so that 0 *≤ q*_*i*_ *≤* 1; see Figure S1a). The quality also influences the maximum amount of help an individual can provide to a partner in need. The maximum help is

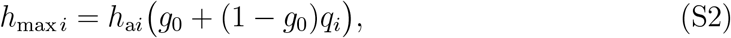

with, for instance, *g*_0_ = 0.75. A top quality individual (*q*_*i*_ = 1) can then donate a maximum amount of *h*_max *i*_ = *h*_a*i*_.

### Bond strength dynamics

For an individual *i* who requests help a given day, i.e. who has *z*_*i*_ = 0, there is updating of the bond strength. We assume that the starting value of bond strength is *x*_s_, e.g., *x*_s_ = 1.25, which should be large enough that at least some help is donated early in a partnership (see Fig. 1c). After a request from *i* to a particular *j, i* updates the value *x*_*ij*_ using the rate *α*_*i*_ and an asymptotic (maximum) bond strength *x*_a_, e.g., *x*_a_ = 20.0. So, with *u*_*ji*_ the amount of help from *j* to *i*, the update is

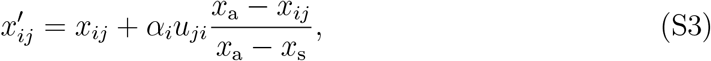

where 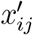 is the updated value of *x*_*ij*_. The donating individual *j* also updates its bond strength *x*_*ji*_, as follows:

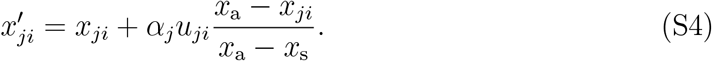

These are the same as equations (1, 2) in the main text. See also Figs. 1a, 3a, S5a.

This bond strength dynamics is inspired by the Rescorla-Wagner (RW) model of classical conditioning. The interpretation is as follows. The asymptotic bond strength *x*_a_ corresponds to *λ* in the RW model, and might be considered a property of the partner, but for simplicity we let it be the same for all partners (we could have higher *x*_a*j*_ for partners *j* of higher quality). The rate *α*_*i*_ corresponds to the RW *α*. The interpretation of the amount of help *u* in equations (S3, S4) is analogous to the RW parameter *β* (or, instead, *u*_*ji*_*/*(*x*_a_*−x*_s_) is analogous to RW *β*). In any case, having *x*_a_*−x*_s_ in the denominator simplifies the interpretation of the dynamics; (*x*_a_ *− x*_*ij*_)*/*(*x*_a_ *− x*_s_) starts out at 1 and then approaches 0 as the bond strength builds up.

### Estimates of association

Each individual *i* maintains an estimate *ρ*_*ij*_ of the association between itself and and another group member *j*. The estimates are updated daily, when individuals are in their chosen places. The update for *i* is

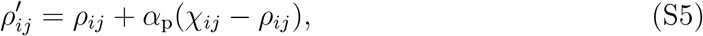

where *χ*_*ij*_ is either one or zero and indicates whether *i* and *j* are in the same place, and *α*_p_ is a place association update parameter (e.g. *α*_p_ = 0.1). The starting value for *ρ*_*ij*_ is assumed to be zero.

### Effective bond strength

The effective strength of the bond between *i* and *j*, as estimated by *i*, is the product of the estimated association *ρ*_*ij*_ between *i* and *j* and *i*’s estimate *x*_*ij*_ of the accumulated exchange of help between *i* and *j*. So *i*’s estimate of the strength of the bond between *i* and *j* is

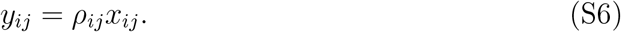

Updating of the association between *i* and *j* is given in equation (S5), and the updating of accumulated helping between *i* to *j* appears in equations (S3, S4).

### Requesting help

If individual *i* failed in foraging, so that *z*_*i*_ = 0, *i* will request help, choosing among available individuals *j*, i.e. those individuals that are in the same place that day. Further, an individual *i* who is in need only considers others that it estimates to have succeeded in foraging, i.e., estimated to have *z*_*j*_ = 1, where the estimate is correct with probability *p*_d_ (e.g., *p*_d_ = 0.98). If there is more than one such alternative, individual *i* uses a soft-max procedure on the effective bond strengths *y*_*ij*_ = *ρ*_*ij*_*x*_*ij*_ to choose the one to ask help from. In addition, individual *i* takes into account if it has previously received help from *j*, or if *j* is a new donor individual. For an established partner, the probability of requesting help from *j* is proportional to

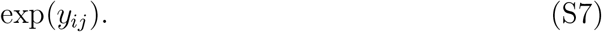

See Figure 1b and equation (3) for an illustration with choice between two individuals. For a new donor *j*, the amount ∆*y*_new *i*_ is added to the effective bond strength, leading to the probability of requesting help from *j* being proportional to

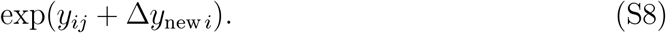

An individual in need can ask help from more than one of the available individuals. In the model, individuals have the opportunity to ask for help twice, from different individuals among the available ones in the current place.

### Group substructure and choice of places

The model allows for the possibility that a group is structured into temporary sub-groups, which we identify with places, e.g., resting places. There are *K* places and each day each group member chooses which place to visit.

We assume that the probability for an individual to select a particular place is influenced by an estimated effective bond strength experienced by the individual in that place, but that there is a small probability *ϵ*_p_ that the individual ends up in a random place (e.g., *ϵ*_p_ = 0.05). Further, to limit the outcome that all individuals in the group select the same place, we assume that there is density dependence in the probability of selecting a place. Let 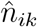 be an estimate by individual *i* of the density (number of individuals) in place *k*, and *ŷ* _*ik*_ an estimate by *i* of the effective bond strength of place *k*. The update rate *α*_p_ is used to update the estimated densities,

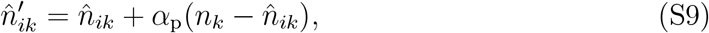

where *k* is the current place for individual *i* and *n*_*k*_ is the actual current density in place *k*. Similarly, the place bond strength is updated as

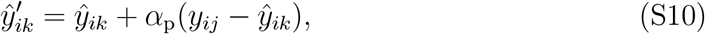

where *y*_*ij*_ = *ρ*_*ij*_*x*_*ij*_ is the effective bond strength to another individual *j* currently in place *k*. The update in this equation is performed for each other individual *j* in the current place.

The probability that individual *i* selects place *k* is then, with probability 1 *− ϵ*_p_, proportional to

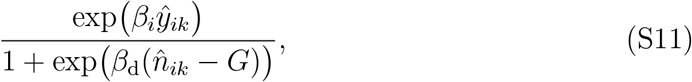

where *β*_*i*_ is the place choosiness trait of *i*, and *β*_d_ is a parameter (e.g., *β*_d_ = 4.0).

The parameter *G* = *N/K* acts as a carrying capacity for place *k*. With probability *ϵ*_p_, however, the choice of place is random. For *K* = 2 and equal density estimates for both places, the result is equation (4) in the main text (see Figure 1b for an illustration).

### Helping amounts

The effective bond strength *y*_*ij*_, as estimated by *i*, is taken into account by *i* when deciding on the amount of help to provide when *j* requests it. In the model, this is expressed as a helping function

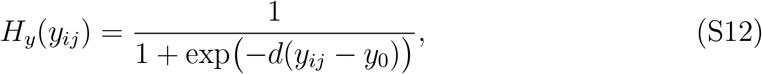

where *d* and *y*_0_ are parameters (e.g., *d* = 8.0 and *y*_0_ = 1.5). The above is the same as equation (5) in the main text. This holds for an established partner *j* that *i* has donated to before, but for a new recipient *j*, the (positive or negative) amount ∆*y*_new *i*_ is added to the effective bond strength *y*_*ij*_, resulting in the value *H*_*y*_(*y*_*ij*_ + ∆*y*_new *i*_), and this is illustrated in Fig. 1c.

The amount of help provided also depends on the total helping balance *w*_*ij*_. The total helping balance between *i* and *j* is the sum of the amounts *u*_*ji*_ of help previously received from *j* minus the sum of the amounts *u*_*ij*_ previously donated by *i* to *j*. Further, let *ζ*_*i*_ denote the current day resource balance for individual *i*. The within-day dynamics of this variable is

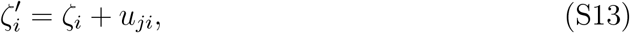

after *i* has received the amount *u*_*ji*_ from *j*, and

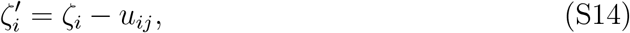

after *i* has donated the amount *u*_*ij*_ to *j*. We assume *ζ*_*i*_ = 0.5*q*_*i*_ at the start of the day. The reason this variable is needed is that an individual *i* might be involved in several exchanges of help on the same day. In the model, the influence of *q*_*i*_, *w*_*ij*_, *z*_*i*_, and *ζ*_*i*_ on the helping amount is expressed as the function

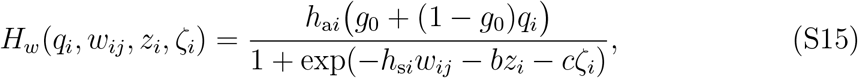

where the parameters *b* and *c* are the same as in the mortality function below (in equation (S17), e.g., *b* = 4.5 and *c* = 3.0). This is the same as equation (6) in the main text. Also note that the numerator in this equation is the value *h*_max *i*_ from equation (S2). See Figure 1d for an illustration of the function. The amount donated from *i* to *j* is the product of the two functions:

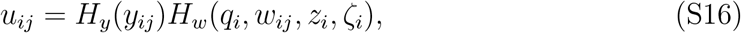

or, for a new recipient, the same but adding the amount ∆*y*_new *i*_ to *y*_*ij*_.

The above means that the social bond mechanism implements a form of state-dependent reciprocity. The reciprocity is expressed in how the amount of help is influenced by the effective bond strength *y*_*ij*_ and by the helping balance *w*_*ij*_. Depending on how ∆*y*_new *i*_ evolves, new individuals can be treated differently from established partners. There is also an influence of the quality *q*_*i*_ on the amount of help an individual provides, as seen from equations (S2) and (S15).

### Survival effects

Survival is the only fitness component acting in the model. The (daily) rate of mortality depends on the foraging success state *z*_*i*_ and the current-day resource balance *ζ*_*i*_. The rate of mortality for an individuals *i* is then

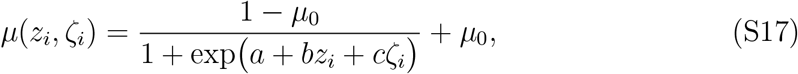

where *a, b, c*, and *µ*_0_ are parameters. This is the same as equation (7) in the main text. We used the parameter values *a* = 2.0, *b* = 4.5, and *c* = 3.0, which means that succeeding in foraging (*z*_*i*_ = 1 vs. *z*_*i*_ = 0) corresponds to receiving an amount of 4.5*/*3 = 1.5 units of help. The parameter *µ*_0_ is a background rate of mortality, irrespective of the individual’s resource state (we used *µ*_0_ = 0.002). The resulting effects on mortality of donating and receiving help are illustrated in Fig. 2a.

In order to evaluate marginal costs and benefits of donating and receiving help, we need the partial derivative of *µ* with respect to *ζ*:

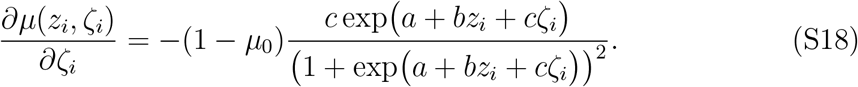

The marginal cost for an individual *i* with *z*_*i*_ = 1 and quality *q*_*i*_, at the point of donating the amount *u*_*ij*_, is minus this partial derivative evaluated at *z*_*i*_ = 1 and *ζ*_*i*_ = 0.5*q*_*i*_ *−u*_*ij*_. The corresponding marginal benefit for an individual *j* with *z*_*j*_ = 0 and quality *q*_*j*_, at the point of receiving the amount *u*_*ij*_, is minus this partial derivative evaluated at *z*_*j*_ = 0 and *ζ*_*j*_ = 0.5*q*_*i*_ + *u*_*ij*_. Using this to examine the marginal benefit-cost ratios for average quality individuals (*q*_*i*_ = 0.5), we find that the ratio for a very small amount is quite large, around 80, whereas at the point of transferring an amount *u* = 0.4, the ratio is 7.9. The ratio then decreases sharply as the amount gets larger, which can be deduced from Fig. 2a.

Individuals that die are replaced through reproduction by surviving individuals. This happens at intervals of *T* days (e.g., *T* = 20). To avoid effects of local relatedness, parents to a new individual are randomly drawn from the global population.

### Model variants

#### Visitors between groups

For this model variant, a number of randomly selected pairs of groups exchange visitors, with results in Fig. S6 and Table S2 (in this case, 20 random pairs out of 250 groups exchanged visitors for a given 20 day interval). For a visiting exchange, each individual in a group had a 50% chance of becoming a visitor in the paired group. We assume that individuals distinguish between visitors and residents, such that individuals have an additional trait ∆*y*_vis,*i*_, with a similar function as ∆*y*_new,*i*_, but applied to visitors (see Table S2). A further possibility, which we did not implement, might be to let some of the visitors stay in the new group, thus becoming residents.

#### No individual recognition

This model variant can be regarded as investigating the evolution of generalized reciprocity, with continuous variation in helping amounts, with results in Fig. S7 and Table S3. We assume that an individual treats all other individuals as the same, with full association (*ρ*_*ij*_ = 1), an already established bond (we used *x*_*ij*_ = 2.5), and a helping balance *w*_*ij*_ formed from all received and donated help. In this way, the helping function *H*_*w*_ in equations (6, S15) and Fig. 1d regulates the exchange of help.

### Details of individual-based simulations

There are two issues that are helpful to keep in mind when performing evolutionary simulations for the model. First, the cost of donating help might be fairly low, which means that selection can be relatively weak. Second, the model has five traits, which also means that the strength of selection on each particular trait can be weak. It is then important for there to be ample and independent amounts of genetic variation in each trait, so that evolutionary change can proceed efficiently. To achieve this in a simple way, we assume that each trait is determined by an unlinked haploid locus, but with sexual reproduction and free recombination, as follows. To produce a haploid offspring, two parents are chosen to form a diploid from copies of their haploid genomes, and the haploid offspring is then formed with recombination and mutation from this diploid (thus, adult individuals can be regarded as gametophytes). Alleles mutate with a fairly high probability of 0.002, and the mutational increments have Laplacian (bi-exponential) distributions (these distributions have long tails), with standard deviations for a locus ranging from 0.10 down to 0.04, chosen so that simulations could readily locate evolutionary equilibria. We use lower and upper limits for allelic (trait) values, with a fairly high upper limit of 8.0 to allow traits to evolve to large values (see Table S1).

Reproduction occurs at intervals of *T* days (*T* = 20), at which time individuals that have died during the interval are replaced by new offspring, with parents randomly selected from the population. Population size for a simulation is at least 4000 (e.g., 175 groups, each with *N* = 24 individuals). For each case reported in Tables S1 to S3, after having reached evolutionary equilibrium, successive simulations were performed over 200, 000 days (approximately 700 generations), repeated at least 100 times, to estimate means and standard deviations of the traits at evolutionary equilibrium.

## References

[1] Leimar O, McNamara JM (2023) Game theory in biology: 50 years and on-wards. Philosophical Transactions of the Royal Society B: Biological Sciences 378(1876):20210509.

[2] Silk JB (2007) The adaptive value of sociality in mammalian groups. Philosophical Transactions of the Royal Society B: Biological Sciences 362(1480):539–559.

[3] Seyfarth RM, Cheney DL (2012) The evolutionary origins of friendship. Annual Review of Psychology 63(1):153–177.

[4] Wilkinson GS (1984) Reciprocal food sharing in the vampire bat. Nature 308(5955):181–184.

[5] Carter GG, Wilkinson GS (2013) Food sharing in vampire bats: reciprocal help predicts donations more than relatedness or harassment. Proceedings of the Royal Society B: Biological Sciences 280(1753):20122573.

[6] Wilkinson GS, Carter GG, Bohn KM, Adams DM (2016) Non-kin cooperation in bats. Philosophical Transactions of the Royal Society B 371:20150095.

[7] Trivers RL (1971) The evolution of reciprocal altruism. The Quarterly Review of Biology 46(1):35–57.

[8] Boyd R, Richerson PJ (1988) The evolution of reciprocity in sizable groups. Journal of Theoretical Biology 132(3):337–356.

[9] Lehmann L, Keller L (2006) The evolution of cooperation and altruism – a general framework and a classification of models. Journal of Evolutionary Biology 19(5):1365–1376.

[10] Connor RC (2010) Cooperation beyond the dyad: on simple models and a complex society. Philosophical Transactions of the Royal Society B: Biological Sciences 365(1553):2687–2697.

[11] Raihani NJ, Bshary R (2011) Resolving the iterated prisoner’s dilemma: theory and reality. Journal of Evolutionary Biology 24(8):1628–1639.

[12] West SA, Cooper GA, Ghoul MB, Griffin AS (2021) Ten recent insights for our understanding of cooperation. Nature Ecology & Evolution 5(4):419–430.

[13] Roberts G, Sherratt TN (1998) Development of cooperative relationships through increasing investment. Nature 394(6689):175–179.

[14] Carter GG, et al. (2020) Development of new food-sharing relationships in vampire bats. Current Biology 30(7):1275–1279.e3.

[15] Aubier TG, Kokko H (2022) Volatile social environments can favour investments in quality over quantity of social relationships. Proceedings of the Royal Society B: Biological Sciences 289(1973):20220281.

[16] Rescorla RA, Wagner AR (1972) A theory of Pavlovian conditioning: Variations in the effectiveness of reinforcement and nonreinforcement. in Classical conditioning II: current research and theory, eds. Black AH, Prokasy WF. (Appleton-Century-Crofts, New York), pp. 64–99.

[17] Barta Z, McNamara JM, Huszar DB, Taborsky M (2011) Cooperation among non-relatives evolves by state-dependent generalized reciprocity. Proceedings of the Royal Society B: Biological Sciences 278(1707):843–848.

[18] Wilkinson GS (1985) The social organization of the common vampire bat. Behavioral Ecology and Sociobiology 17(2):111–121.

[19] Wilkinson GS (1985) The social organization of the common vampire bat. Behavioral Ecology and Sociobiology 17(2):123–134.

[20] Wilkinson GS (1988) Reciprocal altruism in bats and other mammals. Ethology and Sociobiology 9(2):85–100.

[21] Ripperger SP, et al. (2019) Vampire bats that cooperate in the lab maintain their social networks in the wild. Current Biology 29(23):4139–4144.e4.

[22] Ripperger SP, Carter GG (2021) Social foraging in vampire bats is predicted by long-term cooperative relationships. PLOS Biology 19(9):e3001366.

[23] Carter GG, Wilkinson GS (2015) Social benefits of non-kin food sharing by female vampire bats. Proceedings of the Royal Society B: Biological Sciences 282(1819):20152524.

[24] Carter GG, Farine DR, Wilkinson GS (2017) Social bet-hedging in vampire bats. Biology Letters 13(5):20170112.

[25] Carter GG (2021) Co-option and the evolution of food sharing in vampire bats. Ethology 127(10):837–849.

[26] Enquist M, Leimar O (1993) The evolution of cooperation in mobile organisms. Animal Behaviour 45(4):747–757.

[27] Schülke O, Bhagavatula J, Vigilant L, Ostner J (2010) Social bonds enhance reproductive success in male macaques. Current Biology 20(24):2207–2210.

[28] Gerber L, et al. (2020) Affiliation history and age similarity predict alliance formation in adult male bottlenose dolphins. Behavioral Ecology 31(2):361–370.

[29] Schino G, Aureli F (2009) Reciprocal altruism in primates: partner choice, cognition, and emotions in Advances in the Study of Behavior. (Academic Press) Vol. 39, pp. 45–69.

[30] Hammerstein P (2003) Why is reciprocity so rare in social animals? in Genetic and cultural evolution of cooperation, ed. Hammerstein P. (MIT Press, Cambridge, MA), pp. 83–93.

[31] Boyd R, Richerson PJ (1992) Punishment allows the evolution of cooperation (or anything else) in sizable groups. Ethology and Sociobiology 13(3):171–195.

[32] McNamara JM, Barta Z, Fromhage L, Houston AI (2008) The coevolution of choosiness and cooperation. Nature 451(7175):189–192.

[33] McNamara JM, Leimar O (2020) Game theory in biology: Concepts and frontiers. (Oxford University Press, Oxford), First edition.

